# An axonal sodium current gates motoneuron doublets and force amplification

**DOI:** 10.64898/2026.07.29.741588

**Authors:** Robin Rohlén, David Bennett, Monica Gorassini, Dario Farina

**Author notes:** Correspondence: Robin Rohlén and Dario Farina.

## Abstract

Brief high-frequency bursts of action potentials shape neural coding. In spinal motoneurons, they appear as doublets: closely spaced spike pairs, observed for a century, that amplify muscle force nonlinearly through the catch-like property. However, the cellular and anatomical origins of doublets remain unresolved. Combining intracellular recordings, a conductance-based model, and human motor unit and force data, we show that initial and repetitive doublets arise from the same mechanism. Both spikes are initiated at the axon initial segment, but the second is driven by a persistent sodium (NaP) current at the first node of Ranvier, which returns toward the initial segment and sums with somatic NaP current, the calcium-mediated afterdepolarization, and the passive membrane response. In our model, blocking any of these active currents abolishes the doublet, and monoaminergic facilitation of NaP determines whether doublets occur. Human motor units discharged repetitive doublets whose interval lengthened over time, a time course set by inactivation of the somatic NaP current, and produced nonlinear increases in force. Because monoaminergic drive both gates the doublet and sets how its interval evolves, neuromodulation shapes not only the gain but also the timing of motoneuron output.

## Introduction

Brief high-frequency spike bursts are a common pattern across neural systems and can profoundly shape information transfer and output^1–3^. In spinal motoneurons, such bursts occur as closely spaced (<10 ms) spike pairs, or doublets, which were first described nearly a century ago in the cat^4^. Since then, it has been well established that motoneurons can generate doublets during recruitment or reflex-driven activation in cats^5,6^, rats^7^, and humans across a range of tasks^8–13^. Observations from human experimental studies have classically distinguished between initial doublets at recruitment and repetitive doublets during sustained activation near threshold, suggesting different mechanisms underlying the two types^10^. However, this categorization is largely empirical, and the underlying physiology remains poorly understood.

Functionally, doublets nonlinearly enhance muscle force by evoking summating Ca^2+^ transients inside muscle fibers, a phenomenon known as the catch-like property^14–16^, providing an efficient means of boosting early force output. Although such nonlinear force enhancement has not yet been demonstrated at the level of individual human motor units during voluntary contractions, the presence of doublets in healthy humans and their prevalence in conditions such as amyotrophic lateral sclerosis (ALS)^17–20^ highlight their potential relevance for both normal motor control and neuromuscular disease.

Motoneuron doublets are triggered by an afterdepolarization (ADP) following an action potential, which transiently increases motoneuron excitability^21–24^. Early studies showed that blocking Ca^2+^ entry abolished the ADP^25,26^, suggesting that Ca^2+^ currents contribute to this post-spike depolarization. Later work demonstrated that blocking Ca^2+^-activated outward small-conductance potassium (SK) currents unmasks prominent ADPs and promotes high-frequency firing at the doublet interval^27^, indicating that the expression of the ADP depends on the balance between inward depolarizing and outward hyperpolarizing currents.

In motoneurons, persistent inward currents can produce prolonged depolarizations^28^. Subsequent work identified a slowly inactivating persistent sodium (NaP) current capable of sustaining subthreshold depolarization^29^, providing a candidate inward current in addition to the Ca^2+^-mediated ADP to produce doublets. Regarding the anatomical origin that enables prominent ADP and doublets, early interpretations favored dendritic spike invasion and return currents to the soma^30–32^. However, later studies pointed toward more distal axonal regions, including the axon initial segment (AIS)^33^ and the first node of Ranvier^34^, which are specialized for spike initiation and contain high densities of sodium channels^35^.

Evidence from other neuron types supports a role for axonal NaP in burst generation. In cortical neurons, NaP has been linked to rhythmic bursting^36^, and NaP currents at the first node of Ranvier can facilitate bursting^37^. A follow-up commentary on these findings proposed that neurons may operate in distinct bursting modes, facilitated either by axonal or dendritic mechanisms^38^. However, despite nearly a century of observations, it remains unknown whether motoneuron doublets emerge from dendritic, somatic, or axonal mechanisms, and how these mechanisms interact during natural behavior.

Here, we hypothesized that motoneuron doublets emerge from an interaction between these compartments, in which currents from electrotonically distant sites sum within the ADP window at the appropriate time to push the AIS rapidly to threshold for a second spike. In this framework, the soma serves as an integrative hub where axonal and somatodendritic excitability, together with synaptic and neuromodulatory input, jointly determine whether the neuron operates in a regime that permits doublets.

## Results

### Somatic and axonal recordings identify the components of a doublet

To test whether doublets depend on current from electrotonically distant axon compartments that regenerate action potentials, we began by recording from the soma and axon compartments of motoneurons. These recordings provided the empirical constraints for a computational model of doublet formation developed in subsequent sections, and largely recapitulate observations from classical recordings. For our purposes, we defined a doublet as two spikes separated by more than the refractory period but by less than 10 ms, followed by a prolonged inter-spike interval. Intracellular recordings were made ex vivo in the acutely isolated whole adult mouse sacral spinal cord with ventral roots (VRs) attached to ensure that both axons and large dendrites were intact (**Fig. 1a**). In this preparation, we recorded from either the soma or the axon with either antidromic or orthodromic stimulation, performing various manipulations such as adding 5-HT and TTX (**Fig. 1b**).

**Figure 1.**
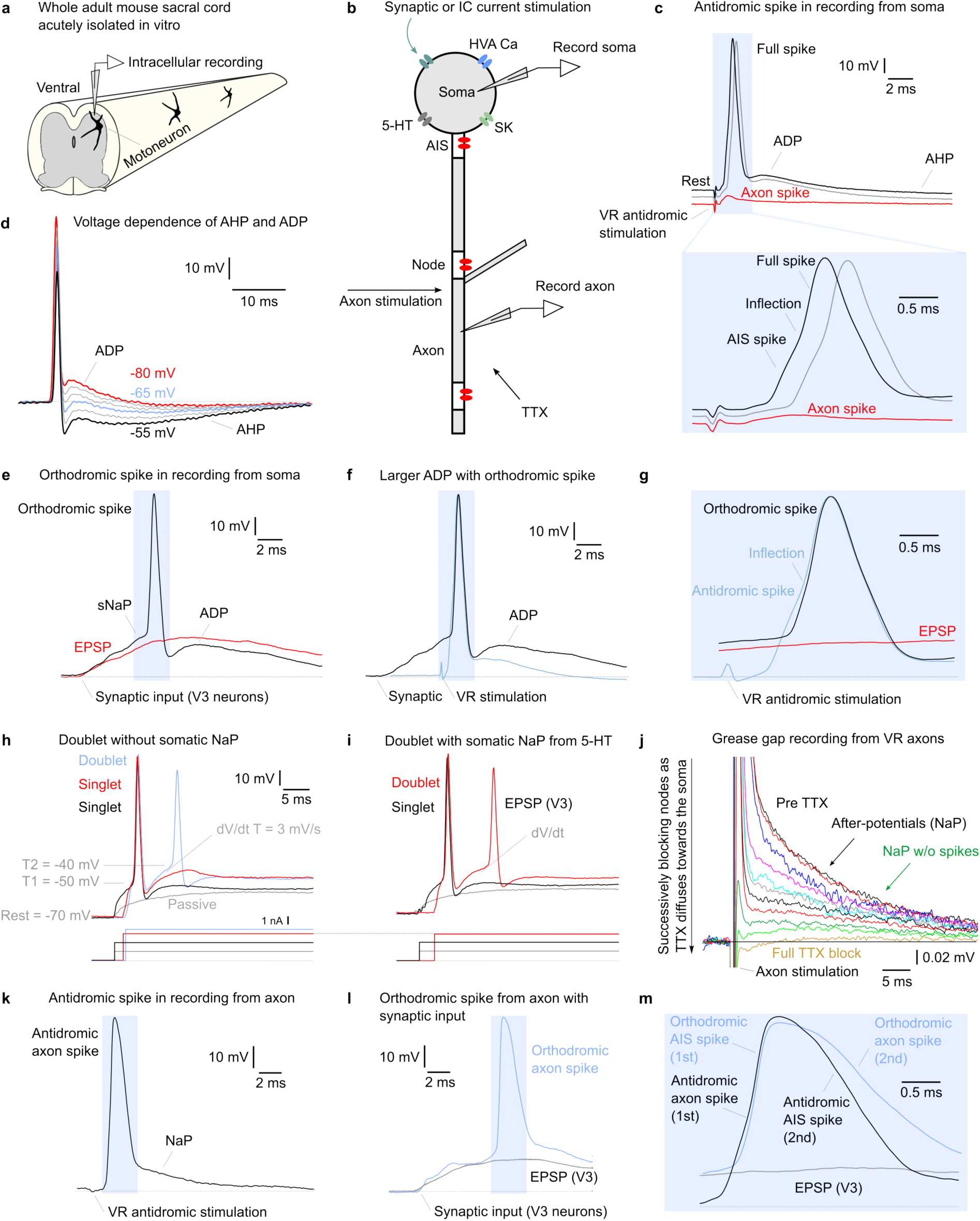
| Somatic and axonal recordings identify the components of a doublet. **a**, The experimental preparation comprises an acutely isolated whole adult mouse sacral spinal cord with attached ventral roots (VRs), preserving intact axons and large dendrites. **b**, Recording configurations: intracellular recordings from the soma or the axon, with antidromic or orthodromic stimulation, and pharmacological manipulations (5-HT, TTX). **c**, Somatic recording during antidromic VR stimulation resolves each compartment in isolation. A small, passively propagated axon spike precedes the axon initial segment (AIS) spike, which in turn triggers the full soma spike after a short delay and inflection. The passively propagated axon spike is seen in isolation when the soma is hyperpolarized to suppress the full spike (red solid line). **d**, The afterdepolarization (ADP), shown in isolation at hyperpolarized potentials near E_K_ (red solid line), where the SK-mediated afterhyperpolarization (AHP) that normally occludes it is abolished. **e**, Orthodromic activation of the soma, the configuration relevant to natural doublet generation. **f**, The ADP evoked by synaptic input (black solid line) is larger than that evoked by VR stimulation (blue solid line). **g**, With orthodromic activation (black solid line), the spike order is reversed relative to antidromic stimulation (blue solid line): the AIS and soma spike occur first, nearly simultaneously and without the AIS inflection, and the axon spike follows. **h**, Response to current injection steps. The passive membrane response rises over an approximately 10 ms time constant (gray solid line), continuing to depolarize through the 10-ms window in which a doublet occurs. Spike threshold is not fixed: at the time of the ADP, the threshold is substantially raised by residual NaV inactivation (-40 mV vs -50 mV, compare T2 with T1). **i**, Without a somatic NaP, a low-current pulse before 5-HT application does not evoke a doublet. After 5-HT restores the NaP, the same step produces a rapid acceleration in membrane potential (dV/dt, lower trace) preceding the doublet. **j**, Grease-gap recording of the composite response of multiple ventral root axons to stimulation at the ventral root entry zone, showing the long afterpotential, which in some axons occurs in the absence of a spike (green traces). **k**, Intracellular recording from a motor axon near the soma during VR stimulation. The antidromic axon spike rises rapidly and is followed not by an AHP but by a long, partly NaP-mediated afterpotential. **l** and **m**, The axon spike evoked by synaptic input (**l**) is broader than that evoked by VR stimulation (**m**).

Antidromic stimulation of the VR axon while recording from the soma revealed a classic sequence of events with evidence of each of the key motoneuron compartments (**Fig. 1c**). A small passively propagated axon spike potential was visible first, as seen in isolation by hyperpolarizing the soma to suppress the full spike. A short time later, the AIS spike arose and quickly triggered the full soma spike, with only a short delay and inflection separating the two. This decomposition of the motoneuron spike into axonal, initial-segment, and somatic components was originally described by Eccles and others^39,40^. For modeling purposes, we consider the AIS and soma-dendritic spikes as a single spike (termed soma spike), since they are electrotonically close.

Following the soma spike, two opposing after-potentials arose whose balance governs whether a second spike and doublet form. The first was a prominent ADP, well known to arise from dendritic and somatic high-voltage-activated (HVA) Ca currents activated by the spike^25,26^. Simultaneous with the Ca current onset, an SK current began and produced a growing after-hyperpolarization (AHP) that occludes the action of the ADP^27^. The ADP was therefore best seen in isolation at hyperpolarized potentials near E_K_, where the AHP goes to zero (**Fig. 1d**). The AHP has two crucial roles during repetitive firing. First, it thoroughly de-inactivates the voltage-gated sodium channels (NaV), allowing the NaP to maximally aid the next spike initiation. Second, it provides a rapidly rising potential at the end of the AHP to trigger the NaV^41^. More generally, NaV spike initiation requires the NaV activation to rise fast enough to escape its own inactivation, so both the rate and the overall level of depolarization are critical^42,43^.

With orthodromic activation of the soma by synaptic input or soma current injection, which is the configuration relevant to natural doublet generation, the same compartments came into play (**Fig. 1e**), but the ADP with synaptic input was larger than the one evoked by VR stimulation (**Fig. 1f**). In addition, the spike order was reversed relative to antidromic stimulation: the AIS and soma spike started first, nearly simultaneously and without the AIS inflection, and the axon spike followed (**Fig. 1g**). Current injection steps revealed passive membrane properties that aid doublet formation, together with spike threshold changes that work against doublets (**Fig. 1h**). That is, a step produced a potential that rose with an approximately 10 ms time constant, so the membrane kept depolarizing through the doublet window and aided initiation of the second spike. Shortly after the first spike, at the time of the ADP, the threshold was substantially higher (T2 vs T1, **Fig. 1h**) due to residual NaV inactivation, making a second spike at this time less likely.

Repetitive firing over seconds, including doublets, usually required a soma NaP, as detailed previously^41,43^. That is, fast-rising steps initiated single spikes but generally did not initiate repetitive firing unless the cell possessed a soma NaP (**Extended Data Fig. 1a-b**). Ramp current injections likewise required a soma NaP, as evidenced by the lack of repetitive firing in motoneurons from the acutely isolated spinal cord; these cells lack a NaP (a linear I-V response), since the brainstem monoamines required for this current were absent (**Extended Data Fig. 1c**). In contrast, facilitating the NaP by adding 5-HT to replace lost brainstem drive enabled repetitive firing, even at long intervals where the AHP had ended (**Extended Data Fig. 1d**). This NaP was quantified during voltage clamp, where it was initiated subthreshold to spiking and, together with the slower-rising Ca^2+^ persistent inward current (Ca PIC), produced a plateau potential that accelerated the membrane toward spike threshold and thereby initiated firing, as detailed previously^41^. Doublets sometimes occurred during these slow ramps, though less frequently than in humans, as detailed later.

The doublet itself depended on this soma NaP-driven acceleration of the potential toward threshold. Without a NaP prior to 5-HT application, the doublet did not occur with low-current pulses, largely because the acceleration in potential was insufficient to trigger a spike within the ADP (**Fig. 1i**). When the NaP was restored by application of 5-HT, the same current step initiated a rapid acceleration in potential prior to the doublet, likely mediated by the somatic NaP (**Fig. 1i**, dV/dt).

When we instead penetrated a motor axon near the soma, its action potential had a different shape, which may aid doublet formation. During VR stimulation, the antidromic axon spike rose very rapidly, unlike in the soma, and was not followed by an AHP but instead by a long afterpotential that allows the axon to aid in generating doublet spikes many ms after the first spike (**Fig. 1k**). The axon spike evoked by synaptic input (**Fig. 1l**) was broader than that evoked by VR stimulation (**Fig. 1m**), presumably because the slow onset of the AIS-soma spike preceded the axon spike, enabling longer-lasting effects that may aid doublet formation. This synaptically evoked axon spike was further elongated by the afterpotential. The long afterpotential was so prominent that it was evident when recording from multiple axons in the ventral root using the grease-gap method (**Fig. 1j**, **Extended Data Fig. 2**), while the axons were stimulated at the ventral root entry zone. Application of low-dose TTX (0.3 µM) slowly blocked the NaP and spikes, progressively reducing the composite action of the afterpotential in discrete steps as TTX diffused to nodes deeper into the spinal cord. Some axons exhibited an afterpotential in the absence of a spike, and this was eventually blocked by TTX (see green traces in **Fig. 1j**). These results indicate that the axon spike afterpotential depends on a very low threshold and long-lasting NaP, consistent with previous reports of strong axonal NaP^44–46^.

Together, these recordings identify the components potentially available to generate a doublet and establish their distinct compartmental origins. The soma and AIS produce a spike and, through spike-evoked Ca currents, a depolarizing ADP that is normally occluded by the SK-mediated AHP. The somatic NaP provides a subthreshold inward current required for repetitive firing and for a fast-rising approach to threshold necessary for a doublet. The axon, in contrast, generates a long, NaP-mediated afterpotential rather than an AHP, offering a depolarizing influence that persists for many milliseconds after the first spike, and is likely central to doublet formation. How these separately identified elements combine in time to produce the second spike of a doublet is the question we address next, using a compartmental model constrained by these recordings.

### A putative temporal sequence for doublet formation

Having identified the components that contribute to action potential formation, we established a putative temporal sequence of events leading to a doublet (**Fig. 2a**), to be investigated in our motoneuron model and in silico experiments. The sequence unfolds over the tens of milliseconds following the first spike, as the separately identified currents come into play in turn and sum to re-trigger the AIS spike.

**Figure 2.**
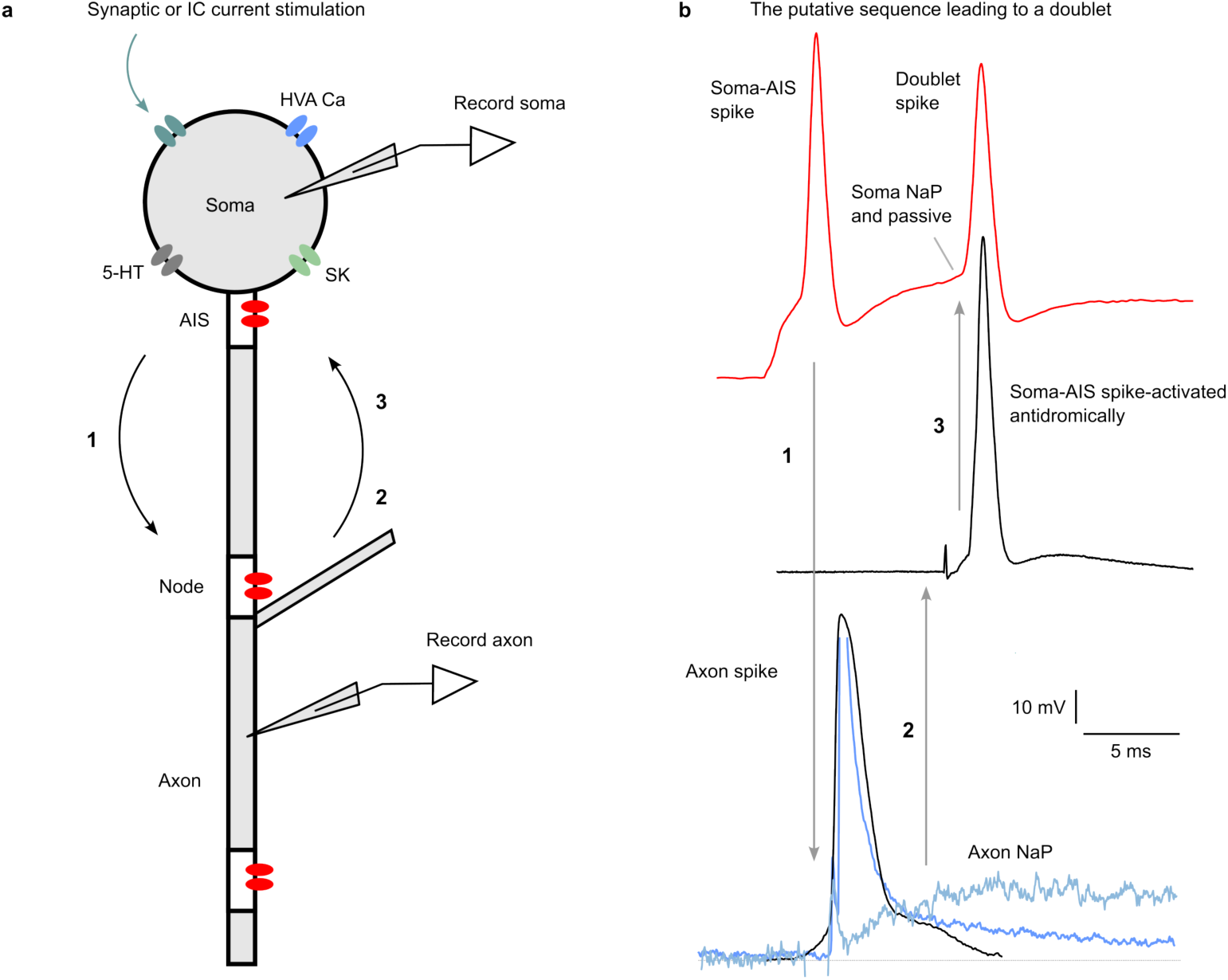
| A putative temporal sequence for doublet formation. **a**, Schematic of the motoneuron compartments and recording configuration. The soma carries high-voltage-activated (HVA) Ca channels, SK channels, and 5-HT receptors, with the electrotonically close axon initial segment (AIS). Sodium channels (red) are concentrated at the AIS and at the nodes of Ranvier. Numbered arrows indicate the proposed sequence: the soma-AIS spike propagates orthodromically to the first node (1); the resulting nodal afterpotential, mediated by an axonal persistent sodium (NaP), returns back toward the AIS (2); and this returning depolarization, summing with the somatic afterdepolarization (ADP) and the recovering somatic NaP, re-triggers the AIS to produce the second spike of the doublet (3). **b**, Recorded potentials corresponding to each step of the sequence, aligned to the schematic in (a). Top (red): somatic recording showing the first soma-AIS spike, the subsequent ADP to which the somatic NaP and the passive membrane rise contribute, and the second (doublet) spike. Middle: a soma-AIS spike activated antidromically. Bottom: the axon spike (black) and the long, NaP-mediated axonal afterpotential (blue), which persists for many milliseconds after the spike. Numbered arrows (1 to 3) link the traces in the order of the proposed sequence.

The putative temporal sequence begins with the first soma spike, which initiates the spike-activated Ca currents that generate the ADP and simultaneously propagate to the axon. For a second spike to follow within the doublet interval, the NaV channels must recover sufficiently from the inactivation produced by the first spike. As shown above, the threshold at the time of the ADP nonetheless remains substantially raised by residual inactivation (T2 vs T1, **Fig. 1h**), so the second spike requires both an adequate ADP and a fast enough approach to this raised threshold. The ADP can, moreover, be masked by the AHP, so the AHP cannot be too large. This link between ADP and AHP has been demonstrated directly by blocking the AHP with apamin^27^: the cell then fires at the doublet interval throughout rather than producing discrete doublets, confirming that the AHP is required for the prolonged interval that separates a doublet from continuous fast firing.

Shortly after the first somatic spike, an axon spike is initiated, with the form shown in **Fig. 2b** when evoked by direct axon activation. Its NaP-mediated afterpotential contributes to the second spike, as confirmed by our simulations (see below). This prolonged axonal afterpotential also extends the net depolarization from the first spike, allowing the AIS NaV channels more time to recover from inactivation before the second spike. In the meantime, the Ca-mediated currents provide the prominent ADP, while the somatic NaP, partially inactivated by the first spike and by the onset of the AHP, recovers and contributes a rising inward current. The sum of these depolarizing currents after the first spike must depolarize the soma rapidly enough to reach the AIS threshold, a process to which the passive rise in potential during a large current step also contributes (**Fig. 2b**, **Fig. 1h-i**).

The relative timing of these currents explains why a doublet requires somatic NaP at low drive levels. As described above, no doublet occurs without a NaP, providing a rapid acceleration in potential preceding the doublet, likely mediated by the somatic NaP (**Fig. 1i**). Although low currents are less likely to produce an initial doublet, several features make them optimal for repetitive doublets. The lower net depolarization reduces the AHP-related masking of the ADP (see **Fig. 1d**), and the lower firing rate allows the AHP to hyperpolarize further and, on its rapid termination, to better trigger the next spike (**Extended Data Fig. 1d**, with AHP depth increasing as firing slows).

Taken together, these observations suggest a putative sequence of events leading to a doublet. First, after the first soma spike, the axon spike occurs and is followed by a long, axon NaP-mediated afterpotential that returns toward the AIS and aids the second spike (**Fig. 2**). In the meantime, the Ca-mediated currents provide a prominent ADP while the soma NaP rises after being partially terminated by the onset of the AHP, a feature that is crucial for the doublet at small current steps. Finally, the axonal and somatic NaP currents sum with the ADP to produce a doublet via recurrent activation of the AIS (**Fig. 2b**). We investigated this sequence in detail using a motoneuron model, as described below.

### A three-compartment model reproduces the putative doublet sequence

To investigate the sequence of events leading to a doublet in detail, we implemented a motoneuron model and conducted in silico experiments providing mechanistic explanations for doublet generation. Such experiments also allowed us to relate the firing patterns to both intracellular recordings and human data. To study the interaction between the dendritic, somatic, and axonal compartments in producing doublets, we implemented a three-compartment conductance-based model (**Fig. 3a**) in which the compartments represent the dendrites, soma, and axon, with the soma lumped with the AIS and the axon compartment representing the first node of Ranvier, matching the compartmental scheme established by the recordings (**Fig. 2a**). The compartments contained 4, 10, and 4 ionic currents, respectively. Fast and slow Na currents (NaF and NaP) were included in the soma and axon (I_NaF,s_, I_NaF,ax_, I_NaP,s_ and I_NaP,ax_). The dendrite and soma included inward calcium currents (the low-voltage-activated I_T_ and the high-voltage-activated I_N_ and I_P_) and an SK current (I_SK_). Model parameters were based on a previous single-compartment model of a neonatal rat hypoglossal motoneuron^47^, adapted to the three-compartment architecture and tuned to match the firing rates observed in our human data (see Methods).

**Figure 3.**
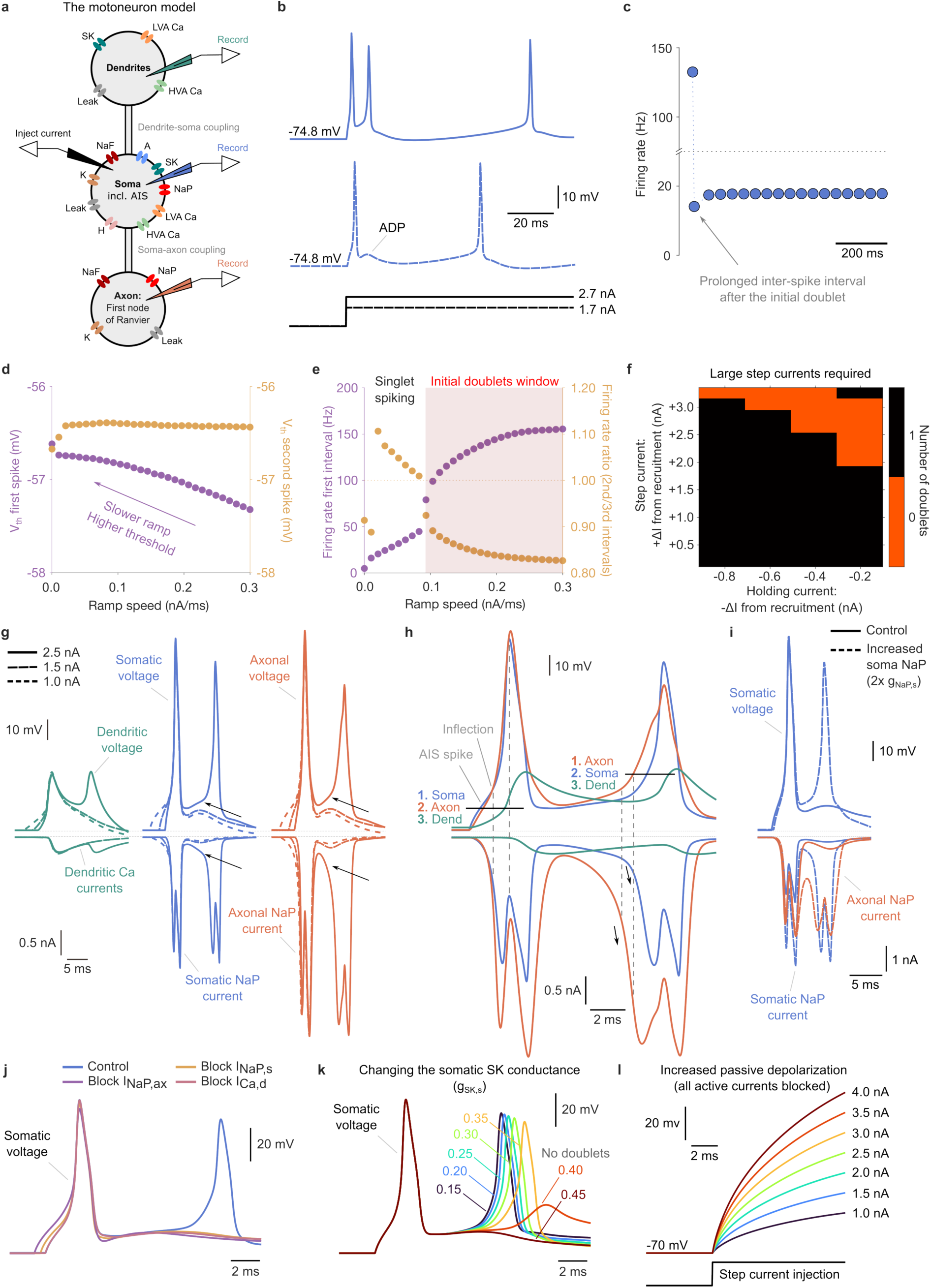
| A three-compartment model reproduces the putative doublet sequence. **a**, Model schematic: The compartments represent the dendrites, the soma (lumped with the axon initial segment, AIS), and the axon (the first node of Ranvier), each containing 4, 10, and 4 ionic currents, respectively. We injected current into the soma and recorded voltage and current from all three compartments and ion channels. **b**, Somatic voltage for two different step currents. A step of 2.7 nA (solid) evoked an initial doublet with spikes approximately 7 ms apart, whereas a step of 1.7 nA (dashed) evoked a single spike followed by a subthreshold afterdepolarization (ADP). **c**, Instantaneous firing rate for the 2.7 nA step. The initial doublet (∼133 Hz) was followed by a prolonged interval (arrow, 14.1 Hz) and then by tonic firing at 17.8 Hz. Note the broken axis. **d**, Voltage threshold (V_th_, the potential at which dV/dt reached 10 mV/ms) of the first spike (purple, left axis) and second spike (gold, right axis) against ramp speed. The first-spike threshold depolarized from -57.3 to -56.6 mV as ramps slowed, whereas the second-spike threshold was independent of ramp speed (∼-56.4 mV). The potential was held at -69.7 mV with a 1.15 nA step current for each ramp. **e**, Firing rate of the first interval (purple, left axis) and the ratio between the firing rates of the second and third intervals (gold, right axis) against ramp speed. The ratio crossed unity (dotted line) at approximately 0.09 nA/ms, separating singlet spiking from the initial doublet window (shaded). We used the same holding and step currents as in (**b**-**c**). **f**, Number of doublets as a function of holding current relative to the recruitment threshold and of step current above it. Black, no doublet; orange, one doublet. Steps of 2.0 to 3.0 nA were required, with smaller steps sufficing at holding currents closer to recruitment. **g**, Voltage (top) and currents (bottom) in the dendritic (green), somatic (blue) and axonal (orange) compartments for three step currents (2.5 nA, solid; 1.5 nA, long dash; 1.0 nA, short dash). A doublet occurred only at 2.5 nA. Arrows mark the voltage rise before the second spike and the accompanying rise in the somatic and axonal NaP currents. The dendrite behaved as a passive load. The potential was held at -74.8 mV. **h**, Overlay of the three compartmental voltages (top) and currents (bottom) around the doublet, at an expanded timescale. The AIS spike appears as an inflection preceding the axon spike. Horizontal lines and vertical dashed lines mark the order of engagement: soma, axon, dendrite at the first spike, and axon, soma, dendrite at the second. Arrows mark the rise in both NaP currents before the second spike. Note the long axonal NaP tail following the first spike. Before the second spike, the axonal compartment depolarizes ahead of the soma, reflecting the earlier activation and slower inactivation of the axonal NaP rather than an earlier spike. The somatic and axonal spikes then occur essentially simultaneously. **i**, Somatic voltage (top) and somatic and axonal NaP currents (bottom) in control (solid) and with the somatic NaP conductance increased twofold (dashed). Increasing g_NaP,s_ converted singlet spiking into an initial doublet, mirroring the effect of 5-HT application (see Fig. 1i). The potential was held at -74.8 mV with a 2.1 nA step current. **j**, Somatic voltage in control (blue) and after blocking the axonal NaP (purple), the somatic NaP (gold) or the dendritic Ca currents (pink). Each block abolished the doublet. The potentials were held at ∼-75 mV with a 2.4 nA step current. **k**, Somatic voltage for somatic SK conductances (g_SK,s_): 0.15-0.45 µS. The intra-doublet interval lengthened with increasing conductance until the doublet was abolished at 0.40 and 0.45 µS. The potentials were held at ∼-75 mV with a 2.7 nA step current. **l**, Somatic voltage with all active currents blocked, for step currents between 1.0 and 4.0 nA, isolating the passive depolarization within the ADP window.

Consistent with our intracellular recordings (see **Fig. 1i**), the amplitude of the step current injected into the soma was decisive for producing an initial doublet, with the two spikes occurring approximately 7 ms apart (**Fig. 3b**). For the smaller current injection, a prominent ADP was evident but was not large enough to trigger a second spike. For the larger step, the doublet, with an instantaneous frequency of ∼133 Hz, was followed by a prolonged inter-spike interval at 14.1 Hz, then tonic firing at 17.8 Hz (**Fig. 3c**).

To investigate whether spike initiation in the model depends on the rate of membrane depolarization, as expected from the accommodation of NaV^48^, we injected current ramps of varying speed and measured the voltage threshold (V_th_) of the first and second spikes (**Fig. 3d**). Threshold was defined as the membrane potential at which dV/dt reached 10 mV/ms^49,50^. The threshold of the first spike depended systematically on ramp speed, depolarizing from -57.3 mV at the fastest ramps to -56.6 mV at the slowest, consistent with sodium channel inactivation accumulating during a slow approach to threshold. In contrast, the threshold of the second spike was essentially independent of ramp speed (approximately -56.4 mV), since the second spike is approached not by the injected ramp but by the intrinsic post-spike trajectory of the AHP and the NaP. This dissociation, whereby only the first spike shows a rate-dependent threshold, matches previous modeling of cat motoneurons^51^, and the first-spike shift is in the same direction as the difference between ramp-evoked and pulse-evoked thresholds measured intracellularly in cat motoneurons (+3.8 ± 3.6 mV)^50^. The relatively hyperpolarized thresholds and the accommodation in the model reflect the NaP, which is active subthreshold and lowers the voltage at which spikes are initiated.

Having established that the threshold of the first spike, but not the second, depends on the rate of membrane depolarization, we next asked how ramp speed determines whether an initial doublet is produced at all (**Fig. 3e**). Increasing the ramp speed progressively increased the firing rate of the first interval, from approximately 5 Hz at the slowest ramps to a maximum of approximately 155 Hz at the fastest ramp. To identify doublets, we used the ratio between the firing rates of the second and third intervals, which falls below unity when the interval following the first pair is prolonged, the defining feature of a doublet. This ratio declined monotonically with ramp speed and crossed unity at approximately 0.09 nA/ms, at which point there was a jump in firing rate of the first interval from approximately 45 Hz to 79 Hz with firing rate ratios of 1.01 and 0.92 (see ‘Initial doublets window’ in **Fig. 3e**). Thus, slower ramps produced only singlet spiking, whereas faster ramps produced an initial doublet followed by a prolonged interval. Initial doublets therefore emerge only above a critical rate of depolarization, consistent with the rate of rise, rather than the membrane potential alone, determining whether the second spike is initiated within the ADP^48^.

To determine which step currents produced doublets, we swept holding and step currents relative to the recruitment threshold. With its baseline parameters (see Methods), the model produced only initial doublets, and only in response to large step currents (**Fig. 3f**). Regardless of the holding current, a step of 2.0 to 3.0 nA was required, and the step needed to produce a doublet was smaller when the holding current was closer to the recruitment threshold. This finding is supported by intracellular current injections in rat motoneurons^52^. The model with these parameters produced no repetitive doublets; these emerged only when E_K_ was raised (from -90 mV to -85 mV), as shown below.

To isolate the mechanism underlying initial doublets, we examined the individual compartments and their currents for three step currents (1.0, 1.5 and 2.5 nA). As the step current increased, ADP increased, and a doublet appeared at 2.5 nA (**Fig. 3g**, blue traces, top). In both the somatic and axonal voltage traces, a pronounced voltage rise occurred immediately before the second spike (**Fig. 3g**, black arrows, top). The lumped dendritic compartment, in contrast, behaved as a passive load, showing only a small voltage deflection. There was little difference in the dendritic Ca currents across the three step currents, whereas the NaP currents in the soma and axon increased markedly before the doublet (**Fig. 3g**, blue and orange traces, bottom). These voltage rises correspond to the accelerating depolarization preceding the doublet seen in the intracellular recordings (see **Fig. 1i**).

To establish the order of events preceding the doublet, we overlaid the membrane potentials and currents of the three compartments (**Fig. 3h**). In the first spike of the doublet, the AIS spike, which was modeled within the soma compartment, appeared first and produced an inflection before the axon spike (**Fig. 3h**, blue trace, top), the same signature seen in the somatic recordings (**Fig. 1c**). The somatic and axonal NaP currents each showed a double peak, associated with the AIS and axon spikes (**Fig. 3h**, blue and orange traces, bottom). The axonal voltage closely followed the somatic voltage, whereas the dendritic voltage rose several milliseconds later (**Fig. 3h**, first horizontal line). Although the dendritic voltage and the Ca-mediated dendritic currents were delayed and declined during the ADP window, they contributed substantially within it (**Fig. 3h**, green traces), whereas the somatic and axonal voltages showed an increasing rate of rise.

Before the second spike, the axonal compartment depolarized ahead of the soma (**Fig. 3h**, second horizontal line). This axonal depolarization did not reflect an earlier spike, since the first somatic and axonal spikes occurred at essentially the same time. This reflected the axonal NaP, where its long tail current after the first spike allowed it to reactivate early, so that it contributed inward current across the ADP window before the somatic NaP did. Both currents then accelerated inward immediately before the second spike, but the axon NaP was well ahead (**Fig. 3h**, black arrows). The axonal depolarization was delivered to the soma through the soma-axon coupling, where it summed with the ADP and the somatic NaP to return the AIS to threshold, as proposed from the intracellular recordings (**Fig. 2**).

In further support of the intracellular recordings, increasing the somatic NaP conductance two-fold caused the model to transition from singlet spiking to an initial doublet (**Fig. 3i**, blue traces, top), mirroring the effect of restoring the NaP with 5-HT in the recordings (**Fig. 1i**). Both the somatic and axonal NaP currents increased within the ADP window, with a steeper rate of rise immediately before the second spike (**Fig. 3i**, blue and orange traces, bottom).

These in silico experiments indicate that both the somatic and the axonal NaP currents are central to doublet generation. However, the ADP itself is known to arise from dendritic and somatic Ca^2+^ currents activated by the spike^25,26^, and dendritic currents have been proposed to contribute to bursting in cortical neurons^38^. To test the respective roles of the somatic NaP, the axonal NaP and the Ca-mediated dendritic currents, we took a control condition that produced an initial doublet (**Fig. 3j**, blue trace) and blocked each in turn. Blocking any one of them abolished the doublet (**Fig. 3j**), showing that all three are necessary and supporting the putative sequence (**Fig. 2**) derived from the intracellular recordings (**Fig. 1**).

Previous recordings have also shown that the ADP is opposed by the AHP, as demonstrated by the effect of apamin^27^. To examine how AHP depth affects the doublet, we varied the somatic SK conductance (g_SK,s_) from 0.15 to 0.45 µS. The intra-doublet interval was shortest at the lowest conductance and lengthened progressively as the conductance increased, until the doublet was abolished (**Fig. 3k**).

Finally, in addition to the currents that generate the ADP (the somatic and dendritic Ca currents), oppose it (the SK current), and provide the rate of rise and the final push above threshold (the somatic and axonal NaP), the intracellular recordings showed that the passive membrane response also contributes within the ADP window (**Fig. 1h-i**). To isolate this contribution in the model, we blocked all active currents and injected step currents of increasing amplitude. Larger steps produced correspondingly larger passive depolarizations within the ADP window (**Fig. 3l**), confirming that the passive rise also adds to the depolarization available to reach threshold for the second spike.

Taken together, the model reproduced the sequence of events inferred from the intracellular recordings (**Figs. 1-2**). An initial doublet requires a sufficiently large and fast-rising input, since the second spike must overcome a threshold raised by residual sodium inactivation and can only do so if the membrane approaches it rapidly enough. Within the ADP window, the depolarization available for the second spike is supplied by the Ca-mediated ADP, the passive membrane response, and the somatic and axonal NaP currents, with the axonal NaP contributing a long tail after the first spike and both NaP currents accelerating immediately before the second. The AHP sets the interval by opposing the ADP, so that increasing the SK conductance lengthens the intra-doublet interval and ultimately abolishes the doublet. Blocking the somatic NaP, the axonal NaP or the dendritic Ca currents each abolishes the doublet, establishing that these components are jointly necessary. The first spike is initiated at the AIS, and the second is led by the first node of Ranvier in the axon, consistent with the proposed sequence from the intracellular recordings (**Fig. 2**), confirming that the doublet arises from the recurrent re-activation of the AIS by currents returning from the axon and summing with the Ca-mediated ADP, the somatic NaP, and the passive current.

### Doublets during rapid human contractions reveal a distinct sub-pool of motoneurons recruited for fast force oscillations

As the three-compartment model reproduced the sequence of events inferred from the intracellular recordings, we also sought to relate the modeling to human data through their firing patterns. To investigate the prevalence of initial doublets in rapid contractions in humans, N = 14 healthy subjects performed a slow isometric force ramp to 10% MVC followed by superimposed fast 2 Hz sinusoidal isometric contractions (5-15% of maximum voluntary contraction, MVC) via a force-tracking ankle dorsiflexion task. Some subjects also performed 1 Hz sinusoidal isometric contractions. These fast, low-force contractions span the range in which most people operate during daily activities^53^, e.g., walking to work or swatting a fly. While performing the task, we recorded 64-channel surface electromyography (EMG) and resulting force production (**Fig. 4a**). We mapped the force to a cursor on a screen, where subjects were instructed to track the predefined path and match the amplitude and frequency of the oscillations as best they could. The recorded EMG signals were separated into motoneuron spike trains using a blind source separation algorithm^54^, enabling sampling of a population of motoneuron spike trains innervating muscle fibers of the tibialis anterior muscle.

**Figure 4.**
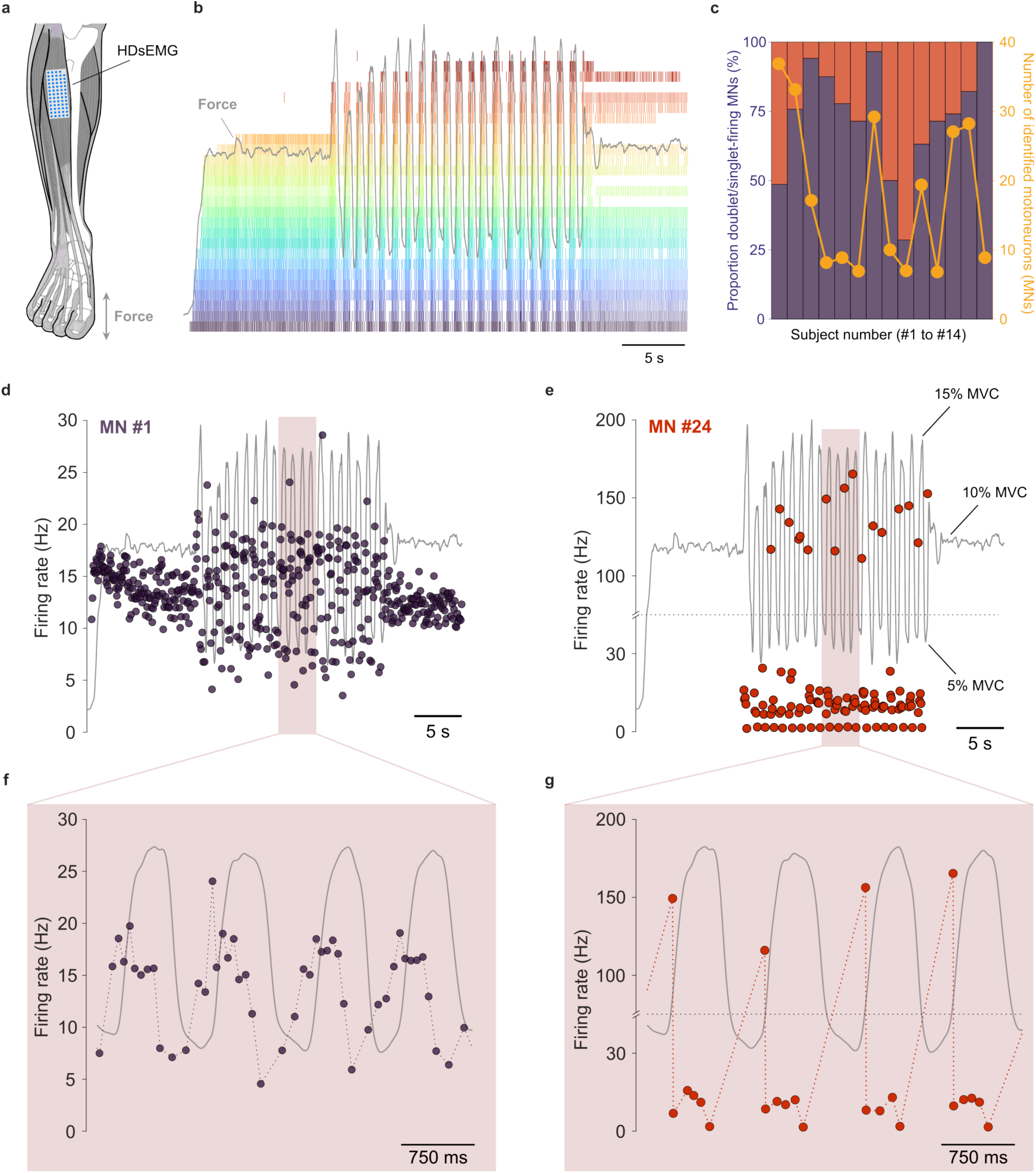
| Doublets during rapid human contractions reveal a distinct sub-pool of motoneurons recruited for fast force oscillations. **a**, Experimental setup and task: healthy subjects (N = 14) performed a sinusoidal ankle dorsiflexion force-tracking task with slow ramp-up, steady force (10% MVC), and fast 2 Hz sinusoidal isometric contractions and occasionally 1 Hz sinusoids, while force and 64-channel surface EMG were recorded. **b**, Representative motoneuron (MN) spike trains in a 1 Hz sinusoidal task from subject #12 showing two distinct activation profiles: a sub-pool recruited early during the ramp phase and a separate sub-pool recruited during the fast oscillations. Doublets occurred predominantly during the sinusoidal phase, with occasional single initial doublets during the ramp. **c**, In N = 14 healthy subjects who performed fast 2 Hz sinusoidal isometric contractions, the proportion of detected motoneurons producing doublets ranged from 29% to 100% across all subjects. Doublets were observed in every subject. **d** and **f**, Firing pattern from MN #1 in (**b**) exhibited a phase-advanced firing pattern relative to the force oscillations. **e** and **g**, Firing pattern from MN #24 in (**b**) produced a doublet preceding each rapid force increase.

During these rapid sinusoidal tasks, we found that a subset of detected motoneurons was activated early in the ramp (**Fig. 4b**). In contrast, another sub-pool was activated later during the fast oscillations superimposed on the ramp (**Fig. 4b**). The detected doublets were mainly activated during the sinusoids in the bursting motoneurons, although single initial doublets occasionally occurred during the ramp phase. All 14 subjects had motoneuron spike trains that produced doublets (**Fig. 4c**). The proportion of detected motoneurons producing doublets ranged from 29% to 100% across all subjects (**Fig. 4c**).

A sub-pool of motoneurons that were activated during the ramp phase exhibited a phase-advanced firing pattern relative to the force oscillations (**Fig. 4d,f**). On the other hand, many of the motoneurons that were activated periodically (bursts) during the fast oscillations produced a doublet before the rapid increase in force (**Fig. 4e,g**). That is, these motoneurons that produced initial doublets did so when they were recruited by fast synaptic input to quickly pass the threshold, similar to the step or fast ramp currents in the in silico modeling (**Fig. 3**). The estimated motoneuron spike trains showed a high signal-to-noise ratio and prolonged inter-spike intervals after each doublet (**Extended Data Fig. 3**). In fact, the periodic bursting patterns resemble the patterns observed from motor units in a human tibialis anterior muscle during imposed muscle stretches while maintaining a constant contraction torque as a motor sinusoidally rotated the ankle^13^. Such a task is similar to walking, in which a previous study reported the presence of doublets^55^. Moreover, different isometric force ramp tasks requiring fast force oscillations also resulted in doublets in the deltoid and triceps muscles in a rhesus macaque^56^, further supporting the prevalence of such bursting patterns across different species, muscles and tasks.

Taken together, these results show that during fast force oscillations at low force levels, as we typically operate daily, a distinct sub-pool of motoneurons is preferentially recruited to produce doublet firing that precedes the rapid force production, consistent with catch-like nonlinear force enhancement during motor tasks. These patterns were highly prevalent among our subjects and consistent with previous studies and our simulations, suggesting that they are common during everyday movements.

### Repetitive doublets share the initial doublet mechanism and adapt through somatic NaP inactivation

In our in silico experiments described above, our motoneuron model produced only initial doublets with a large step current (**Fig. 3f**), likely because the outward drive between spikes via the AHP was too strong to permit repetitive doublets. By reducing the outward drive by changing the potassium reversal potential (E_K_) from -90 to -85 mV in the three-compartment model (dendritic, somatic, and axonal compartments; **Fig. 5a**), the model produced repetitive doublets in the somatic compartment. For example, when a holding current set the resting potential to -79.4 mV and a 0.6 nA step current (just above recruitment threshold) was injected (**Fig. 5b**, blue solid line), the dendritic and axonal compartments showed prominent depolarizations during the post-spike ADP window (**Fig. 5b**, green and orange solid lines). The intra-doublet interval lengthened across the train, so that the intra-doublet rate declined from ∼180 Hz initially to ∼150 Hz after about 1 second, with an inter-doublet rate of ∼6 Hz (**Fig. 5c**). This doublet-frequency adaptation pattern also occurred in healthy humans (see **Fig. 6** below). By varying the soma-axon coupling conductance, we found that repetitive doublets occurred only within a narrow range of coupling (0.19 to 0.20 µS). Weaker coupling produced an initial doublet followed by single spiking (0.13 to 0.17 µS), and the weakest and strongest couplings produced single spiking throughout (0.10 to 0.12 and 0.21 to 0.25 µS; **Fig. 5d**). This dependence on coupling strength supports the axonal contribution to doublet generation, since the coupling sets how effectively the axonal afterpotential returns to the initial segment.

**Figure 5.**
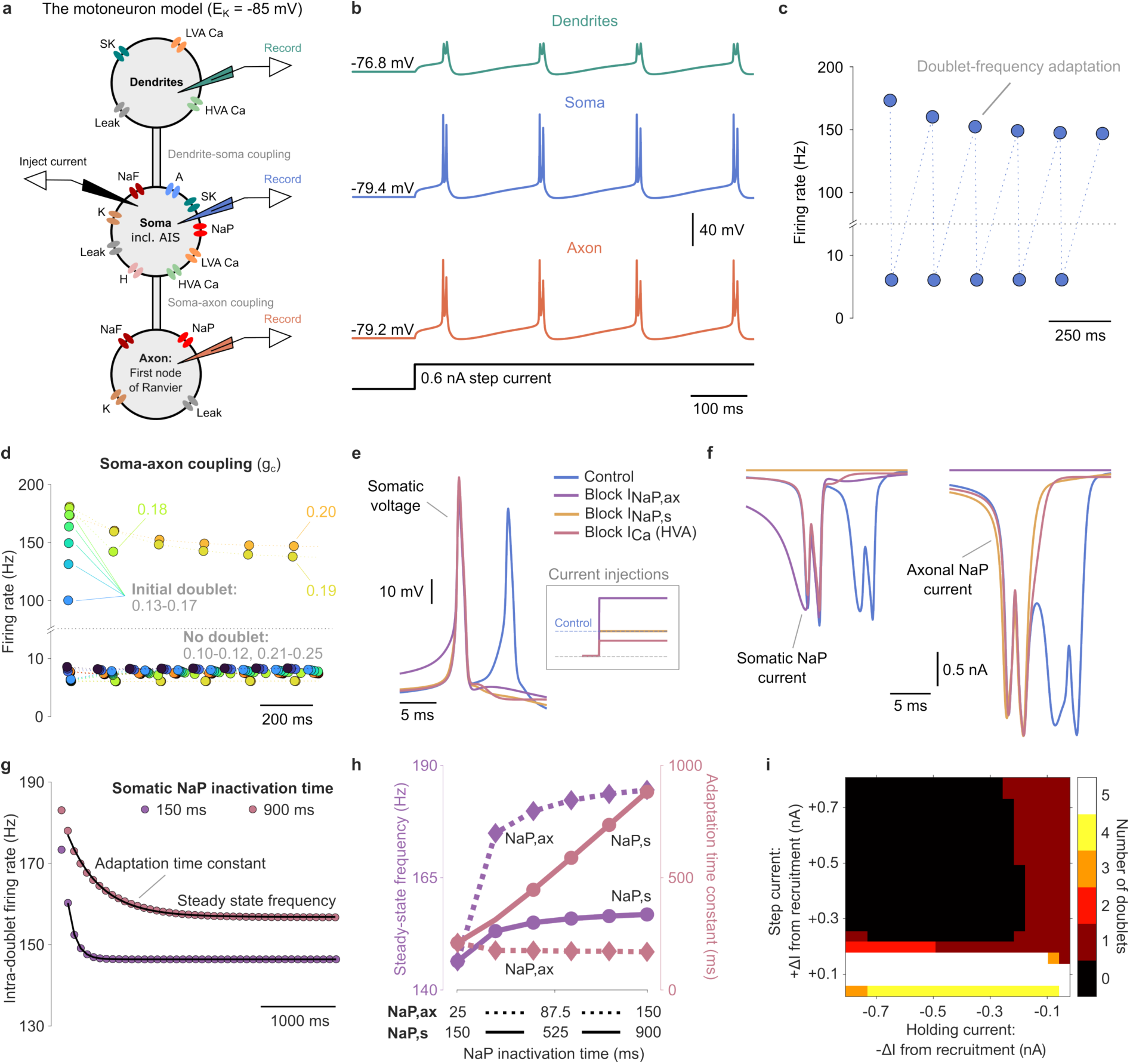
| Repetitive doublets share the initial doublet mechanism and adapt through somatic NaP inactivation. **a**, Schematic of the three-compartment model (dendrite, soma including the AIS, axon), as in Fig. 3a. The potassium reversal potential (E_K_) was raised from -90 to -85 mV, reducing the outward drive between spikes. **b**, Somatic (blue), dendritic (green) and axonal (orange) voltage during repetitive doublets. A holding current set the resting potential to -79.4 mV, and a 0.6 nA step, just above recruitment threshold, was injected into the soma. The dendritic and axonal compartments show prominent depolarizations within the post-spike ADP window. **c**, Instantaneous firing rate for the trace in (**b**). The intra-doublet rate declined from approximately 180 Hz to approximately 150 Hz over the first second, whereas the inter-doublet rate was approximately 6 Hz. Note the broken axis. **d**, Instantaneous firing rate for a range of soma-axon coupling conductances (g_c,sa_, 0.10 to 0.25 µS). Repetitive doublets occurred only within a narrow range of coupling (0.19 to 0.20 µS). Weaker coupling produced an initial doublet followed by single spiking (0.13 to 0.17 µS), and the weakest and strongest couplings produced single spiking throughout (0.10 to 0.12 and 0.21 to 0.25 µS). We used the same holding and step currents as in (**b**-**c**). **e**, Somatic voltage in control (blue) and after blocking the axonal NaP, the somatic NaP or the somatic and dendritic HVA Ca currents. Each block abolished the repetitive doublets. We adjusted the holding currents (∼-73 mV) and step currents (0.300, 0.300, 0.700, 0.185 nA) in each condition to match firing rates (inset). **f**, Somatic and axonal NaP currents associated with the doublet in (**e**). Both accelerate inward immediately before the second spike. The axonal NaP inactivates more slowly and carries a larger current across the ADP window. **g**, Doublet-frequency adaptation during a 10 s step (0.6 nA, holding potential -79.4 mV) for somatic NaP inactivation time constants ranging from 150 to 900 ms, with exponential fits (150 ms is the control for the soma, 25 ms for the axon, see Methods). The first intra-doublet interval was excluded from the fits, since the first doublet is produced by the step transition rather than by the steady input. **h**, Steady-state intra-doublet frequency (left y-axis) and adaptation time constant (right y-axis) against the somatic and axonal NaP inactivation time constant (x-axis: 150 to 900 ms and 25 to 150 ms, respectively). The steady-state frequency saturated at longer inactivation times for both compartments, whereas the adaptation time constant increased approximately linearly for the somatic compartment but remained constant for the axon. We used the same holding and step currents as in (**b**-**c**). **i**, Number of doublets as a function of holding current and step current, both relative to recruitment threshold. Repetitive doublets occurred across a wide range of holding currents but only within a narrow band of step currents, approximately 0.1-0.2 nA above recruitment.

**Figure 6.**
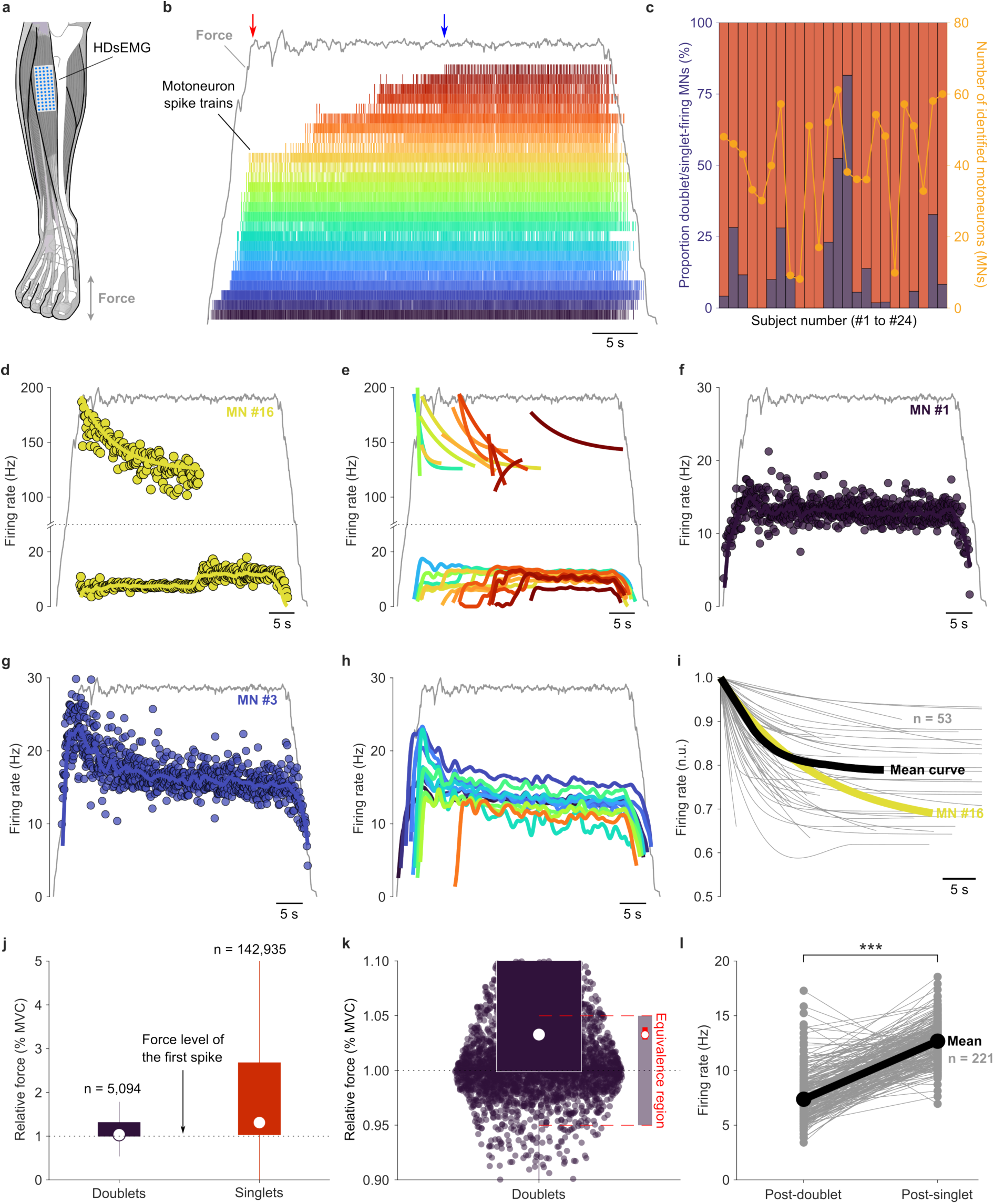
| Repetitive doublets in human motoneurons show doublet-frequency adaptation. **a**, Experimental setup and task: healthy subjects (N = 24) performed a trapezoidal ankle dorsiflexion force-tracking task with slow ramp-up, steady force (2-25% MVC), and ramp-down, while force and 64-channel surface EMG were recorded. **b**, Example of detected motoneuron spike trains from subject #13 during a 20% MVC task with force in gray. **c**, Population summary showing the number of detected motoneuron spike trains and the presence of doublets across subjects. **d**, Representative firing profile from a motoneuron (#16) producing repetitive doublets that transition into singlet spiking. **e**, All repetitive doublet spiking motoneuron profiles from (**b**). **f**, Representative firing profile from a motoneuron (#1) producing singlets correlated with the force. **g**, Representative firing profile from a motoneuron (#3) exhibiting spike-frequency adaptation. **h**, All singlet-spiking motoneuron profiles from (**b**). **i**, Normalized population of doublet-frequency adaptation in repetitive doublets (n = 53 fitted and converged profiles in gray, average in black), showing an average ∼20% decrease in firing rate relative to the initial doublet. **j**, Relative force level of repetitive doublet spikes (5,094 doublet spike pairs vs. 142,935 singlet spikes) compared to the force at the first doublet occurrence, demonstrating clustering near the recruitment threshold, unlike singlet spiking motoneurons. **k**, Equivalence testing (bootstrap TOST) confirms that repetitive doublets occur at force levels statistically equivalent to the recruitment threshold. The shaded horizontal band represents the equivalence region around 1 ([0.95, 1.05]). Black points indicate group medians with 90% bootstrap confidence intervals. **l**, Comparison of effective firing rates during singlet spiking and repetitive doublet spiking, showing significantly lower firing rates during doublet activity. Firing rates above 20 Hz were removed for visual clarity, but statistical analysis was performed on all data points (n = 221 vs n = 230 repetitively doublet-firing motoneurons).

To determine the ionic currents underlying repetitive doublets, we considered a control case producing repetitive doublets (**Fig. 5b**, **Fig. 5e**, blue solid line) and selectively blocked axonal NaP, somatic NaP, and somatic and dendritic HVA Ca currents. As in the initial doublet case, we also found that blocking any of these currents abolished the repetitive doublets (**Fig. 5e**). Note that we varied the step currents to match the firing rates (**Fig. 5e**, inset). As previously shown in the initial doublet case, the axonal and somatic NaP currents exhibited a pronounced inward acceleration just before the doublet (**Fig. 5f**). However, the axonal NaP inactivates more slowly, contributing to a large current within the ADP window.

We examined the doublet-frequency adaptation, a phenomenon that was also clearly observed in our human data (see **Fig. 6**). We injected 10 s step currents (0.6 nA) from a holding current setting the resting potential to -79.4 mV, while varying the somatic NaP inactivation time constant (150 ms as the baseline, see Methods), and fitted an exponential curve to the intra-doublet frequency to estimate the steady-state frequency and adaptation time constant. We excluded the first intra-doublet interval from the fit because the first doublet originates from the holding-to-step current. Somatic NaP inactivation time affected both the steady-state intra-doublet frequency and the adaptation time constant (**Fig. 5g**). Increasing the somatic NaP inactivation time constant from 150 to 900 ms raised both quantities monotonically; the steady-state frequency saturated at longer inactivation times, whereas the adaptation time constant increased approximately linearly (**Fig. 5h**). In contrast, increasing the axonal NaP inactivation time constant from 25 to 150 ms (25 ms as the baseline, see Methods) did only result in a shift of the doublet-frequency adaptation curve to higher frequencies, i.e., no change in adaptation time constant but increased steady-state frequency that saturated at longer inactivation times (**Fig. 5h**). Although a ∼1 s inactivation time for the somatic NaP may sound long, it is indirectly supported by the shape of the doublet-frequency adaptation curve, which is similar to human experimental data (see **Fig. 6d-e**), and by intracellular motoneuron recordings from a chronic spinal rat^57^. This finding suggests that the doublet-frequency adaptation could serve as an experimentally accessible readout of an otherwise hard-to-measure intrinsic property (see Discussion).

In the human literature, repetitive doublets emerge specifically near the recruitment threshold^10^. Our model reproduced this behavior: repetitive doublets occurred across a wide range of holding currents but only within a narrow band of step currents just above recruitment (∼0.1-0.2 nA above threshold; **Fig. 5i**), while larger steps produced singlets unless the holding current was very close to recruitment, when an initial doublet was produced. This near-recruitment preference may seem counterintuitive, since the AHP is larger at low firing rates. However, the depolarizing contributions that drive the second spike are proportionally strongest at low, near-threshold membrane potentials, where the NaP currents are most influential and least inactivated, and the long inter-spike interval allows the slowly inactivating NaP to recover and contribute its returning current within the ADP window. Whether a second spike occurs therefore depends on the margin between this summed depolarization and the AHP, which is most favorable near recruitment, rather than on the size of the AHP alone.

Taken together, initial and repetitive doublets arise from the same mechanism: within the ADP window, the depolarization available for the second spike is supplied by the Ca-mediated ADP, the passive membrane response, and the somatic and axonal NaP currents, with the axonal NaP contributing a long tail after the first spike and both NaP currents accelerating immediately before the second. Opposing these, the SK current sets the AHP, so that a second spike occurs only when the summed depolarization overcomes the AHP within the window. A blockade of any of the depolarizing currents, or a sufficiently strong AHP, abolishes the doublet. Initial and repetitive doublets differ only in how this balance is met: by a strong, fast input that clears the AHP once for initial doublets, and near recruitment, where the balance is reached on successive cycles, for repetitive doublets. The doublet-frequency adaptation was linked to the somatic NaP inactivation, which gradually erodes the depolarizing side across a train, serving as an experimentally accessible readout during natural behavior of an otherwise hard-to-measure intrinsic property.

### Repetitive doublets in human motoneurons show doublet-frequency adaptation

To investigate the properties of repetitive doublets in human motoneurons, N = 24 healthy subjects (14 and 10 subjects from experiment 1 and 2, see Methods) performed a trapezoidal task with a slow ramp followed by a steady force (ranging from 2% to 25% MVC) through a force-tracking ankle dorsiflexion task and ramped down to 0% of the MVC level (**Fig. 6a-b**). As in the sinusoidal task, 64-channel surface EMG and force production were recorded, with force mapped to a cursor on the screen. The EMG signals were then separated into motoneuron spike trains offline.

From the EMG signals, a sub-pool of motoneuron spike trains was extracted, showing different recruitment thresholds (**Fig. 6b**). In this example, from a subject exerting force at 20% MVC, 26 motoneurons were detected, where the first 17 were active when the 20% MVC level was first reached (**Fig. 6b**, red vertical arrow). Over the following 20 seconds, the remaining 9 detected motoneurons were activated, while the force was maintained (**Fig. 6b**, blue vertical arrow).

Many of the detected motoneurons exhibited repetitive doublets, such as motoneuron #16 (**Fig. 6d**), where the estimated spike trains showed a high signal-to-noise ratio and prolonged inter-spike intervals after the repetitive doublets (**Extended Data Fig. 4**). These repetitive doublets exhibited doublet-frequency adaptation, reflecting findings from the in silico experiments and somatic NaP inactivation (**Fig. 5g,h**). Interestingly, the doublet-spiking firing typically transitioned into singlet spiking. In contrast, the interval between doublets did not show such adaptation and was constant, similar to the steady force trace (**Fig. 6e**). In some subjects, we observed doublets intermingled throughout parts of the contraction or the entire contraction, and occasionally multiple doublets in succession (**Extended Data Fig. 5**). Despite their different firing pattern, they also exhibited doublet-frequency adaptation (**Extended Data Fig. 5b,c**).

Another interesting observation was that the motoneurons that repeatedly produced doublets were recruited at different time points during the contraction, even though force remained steady (**Fig. 6e**). Once the repetitive doublets transitioned into single spiking, another motoneuron was typically recruited, producing either doublets or singlets. Notably, the inter-doublet discharge rate (**Fig. 6e**) and the rates of lower-threshold singlet-firing tonic units were stable or declining (spike-frequency adaptation) across the contraction (**Fig. 6f-h**), arguing against a substantial increase in common drive despite this progressive recruitment (see Discussion).

At the population level, of the 24 healthy human subjects who performed the steady force tasks, 16 subjects showed spike trains with at least two doublets in the same recording (**Fig. 6c**). The number of detected motor units was based on all unique motor units from the different trapezoidal force levels (see Methods). A few spike trains contained a single doublet during the ramp phase, but we did not consider these repetitive doublets (at least two were required). On the other hand, no doublets were detected in eight subjects; three of these subjects had a low number of detected motoneurons, whereas five had a moderate-to-high number of detected motoneurons (**Fig. 6c**). In ten subjects, at least 10% of the detected motoneurons produced doublets, whereas in five subjects, at least 25% of the detected motoneurons produced doublets (**Fig. 6c**, purple bars). The number of detected units producing at least one doublet per trapezoidal recording was 3.1 ± 4.6 (min 0; max 22), i.e., 16.7 ± 23.6% (0; 91.7%). By considering only units with at least two doublets, we found 2.1 ± 3.7 (0; 19) units per recording, corresponding to 11.1 ± 19.9% (0; 83.3%).

Although we conservatively defined a doublet as two spikes separated by more than the refractory period but within 10 ms and followed by a prolonged inter-spike interval, we also observed spike pairs separated by slightly more than 10 ms (<100 Hz; for example, 11 to 14 ms), likewise followed by a prolonged inter-spike interval (**Extended Data Fig. 6**). These pairs meet an earlier, more permissive definition, in which a doublet is two spikes less than 20 ms apart (>50 Hz)^58^. Because our criterion is stricter, the number of doublets reported here should be regarded as a lower bound.

We fitted an exponential curve to doublet-frequency adaptation across all motoneurons and subjects, retaining only fits that converged (n = 53). By translating the profiles to the same time point (t = 0) and normalizing to the initial firing rate, we found that, on average, the intra-doublet firing rate decreased by approximately 20% relative to the initial firing rate, with individual adaptation ranging from 5% to 40% (**Fig. 6i**). This adaptation value can be compared to motoneuron #16 (**Fig. 6d**), where the firing rate decreased by approximately 30%.

When previously active doublet-spiking motoneurons transitioned to single-spiking, newly recruited doublet-spiking motoneurons became active while force remained stable (**Fig. 6e**). Repetitive doublets occurred at force levels close to that of the first doublet (1 denotes identical % MVC), whereas singlet-spiking neurons exhibited a broader distribution of force levels relative to the first spike (**Fig. 6j**). Equivalence to 1 was assessed using a nonparametric bootstrap two one-sided tests (TOST) procedure applied to the median (10,000 resamples), with equivalence bounds set to [0.95, 1.05]. The corresponding 90% bootstrap confidence interval [1.028, 1.040] lay entirely within the equivalence region, demonstrating statistical equivalence to 1 (**Fig. 6k**). These results indicate that repetitive doublets occur near the recruitment threshold, consistent with simulations (**Fig. 5i**) and previous work^10^.

Some subjects also performed triangular contractions, i.e., from 0% MVC to 30% MVC over a slow ramp of 15 seconds, before reducing the force over the following 15 seconds down to 0% again (**Extended Data Fig. 7a**). We also observed repetitive doublets during these contractions, but they occurred only within a narrow force range during the ramp (**Extended Data Fig. 7b**). The intra-doublet intervals also increased in these units, showing a doublet-frequency adaptation. We observed one motoneuron that produced only doublets, but this unit was recruited close to the 30% MVC level (see MN #14 in **Extended Data Fig. 7b**). These findings provide additional evidence, with a different target type, that repetitive doublets in healthy subjects are produced close to the motoneurons’ recruitment threshold and show doublet-frequency adaptation.

The doublets also exhibited inter-doublet rates (between the repetitive doublets, i.e., the time between the second spike of a doublet and the first spike of the next doublet) that were considerably lower than for singlet spiking (**Fig. 6d-e**). This difference relates to the summation of AHPs^31^. During the tonic 20% MVC contractions, the median firing rate during singlet spiking was 12.7 Hz, and the inter-doublet rate was 7.4 Hz (**Fig. 6l**). The firing rate of the singlet spiking was approximately 1.7 times that of the doublet spiking. The paired median difference of singlet and doublet firing rates was significantly greater than zero (Wilcoxon signed-rank test: n = 230, W = 1025, p < 0.001).

Taken together, these results demonstrate that repetitive doublet firing is a motoneuron firing pattern present during steady low-to-moderate force contractions in many human subjects, characterized by recruitment at threshold, distinct doublet-frequency adaptation of the doublet discharges, and lower firing rates than singlet spiking, suggesting repetitive doublets as a structured and physiologically distinct firing mode rather than a rare or incidental phenomenon.

### Repetitive motoneuron doublets amplify twitch responses

Functionally, initial doublets nonlinearly enhance muscle force^14–16^. Such nonlinear force enhancement has not yet been demonstrated at the level of individual human motor units during voluntary contractions, and, more importantly, the functional output of repetitive doublet firing remains unclear. Although our results from the human experiments of repetitive doublets indicate force-enhancement capabilities (see the red and blue arrows in **Fig. 6b**), we investigated the force-enhancement capabilities in detail by quantifying the force-related response (FRR) of muscle fibers during repetitive doublets and singlet spikes from the same motor unit.

In a subset of subjects performing trapezoidal contractions, we recorded ultrafast ultrasound from the tibialis anterior muscle, with the probe placed between two electrode grids to capture cross-sectional activity perpendicular to the muscle fibers (**Fig. 7a**). The ultrasound yielded localized intramuscular velocities at a kHz rate^59^, reflecting the thickening of muscle fibers during contraction. From these velocities, we extracted FRR of muscle fibers innervated by individual motoneurons within a region of interest (ROI)^60^ and derived contractile parameters from the FRR^61–63^.

**Figure 7.**
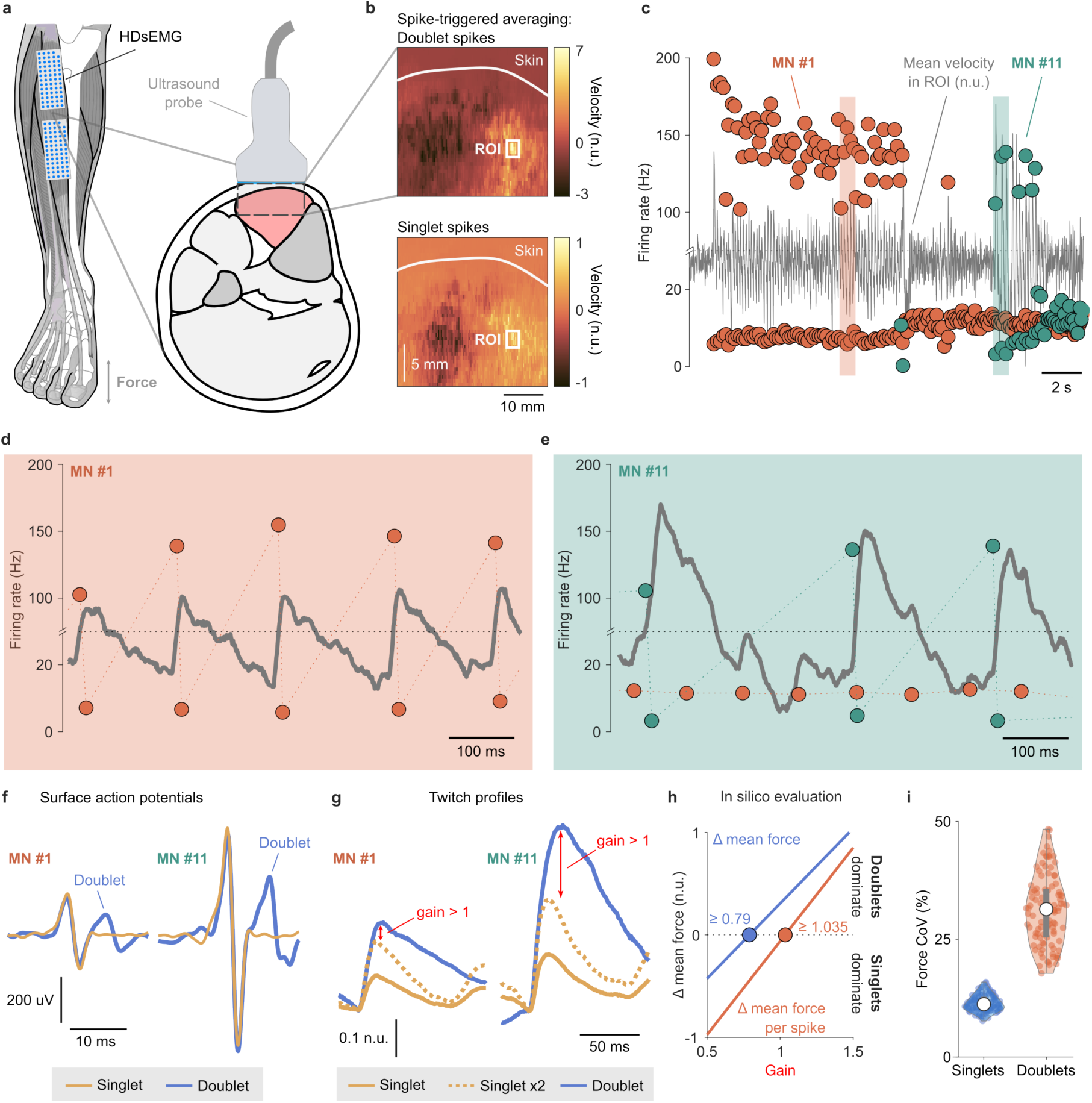
| Repetitive motoneuron doublets amplify twitch responses. **a**, Experimental setup for ultrafast ultrasound imaging of the tibialis anterior, positioned between two surface EMG grids that record cross-sectional activity. **b**, Force-related response (FRR) maps from subject #24 obtained by spike-triggered averaging (by singlet spikes or by the first spike of repetitive doublets) of ultrasound phase shifts along the depth axis for two motoneurons (MN #1 and MN #11). **c**, Average FRR traces extracted from the region of interest (ROI) in (**b**). **d** and **e**, Doublet discharges were phase-advanced to a prominent rise in FRR for both motoneurons, consistent with observations during fast sinusoidal contractions. **f**, Surface motor unit action potentials triggered by singlet and doublet spikes, showing a conserved first peak and a distinct second peak within the doublet. **g**, Average FRR twitch profiles: doublet firing produced twitch amplitudes exceeding twice that of singlet firing (gain > 1). **h**, Simulations of an unfused tetanus from a single motor unit showing the difference in mean force (blue solid line) and force per spike (orange solid line) between doublet and singlet firing as a function of twitch gain. Circles indicate gain values at which the difference crosses zero. **i**, The force variability in terms of the coefficient of variation (CoV) of the simulated singlet and doublet force traces from (**h**) with gain = 1.035, keeping the middle 8 seconds.

We used the spikes from a motoneuron producing repetitive doublets to trigger the FRR and produce two separate FRR maps (**Fig. 7b**), one based solely on triggers from the first spike of a doublet and the other only from singlet spikes (**Fig. 7b**). From the ROI of the FRR maps, we computed the average FRR inside the ROI over time, demonstrating enhanced FRR upon activation of two different motoneurons (MN #1 and MN #11) during their repetitive doublets (**Fig. 7c**). Each doublet was phase-advanced relative to a prominent rise in velocity for both MN #1 (**Fig. 7d**) and MN #11 (**Fig. 7e**), similar to what we observed in the fast sinusoidal contractions (**Fig. 4g**). From the spikes as triggers, we computed the motor unit action potentials at the surface of the skin from one of the electrodes, showing that the first peak of the doublet coincided with the singlet, whereas the second peak of the doublet was different from the first (**Fig. 7f**), which has been reported previously in signals from intramuscular electrodes^64^.

The average doublet twitch was more than twice the amplitude of the singlet twitch, resulting in a gain above one for both MN #1 and MN #11 (**Fig. 7g**). These human recordings show that doublets occur repeatedly during voluntary contraction and produce nonlinear force enhancement, so that, like initial doublets, repetitive doublets also amplify force.

Using simulations of an unfused tetanus from a single motor unit with stochastic firing and a twitch model^65^, we examined how amplification of twitch force during doublets influences force output relative to singlet spiking. Spike trains were generated over 10 s with singlet firing rates drawn from 11.2 to 13.6 Hz and doublet rates set to 56 to 75% of the singlet rate, matching the ranges observed in our human motor unit recordings (**Fig. 6l**). Twitch contraction times were sampled from 75 to 90 ms, and inter-spike intervals were jittered with a coefficient of variation of 10%, yielding 100 stochastic realizations per condition. We considered gain values ranging from 0.5 to 1.5, where the gain is a multiplier of the sum of twitch forces of the doublet (see **Fig. 7g**). Across this physiologically plausible parameter space, increasing the gain applied to twitch amplitude during doublets progressively shifted the balance of force production on the single motor unit level. When the mean force was considered, doublet firing produced a lower force than singlet firing at low gain values but exceeded the singlet force beyond a gain of 0.79 (**Fig. 7h**, blue solid line). In other words, the twitch force in response to a doublet only had to be 1.6 times the twitch force in response to a singlet to produce the same mean force.

When force was instead normalized to the number of spikes, serving as a proxy for energetic cost, the crossover occurred at a higher gain (approximately 1.035, i.e., 2.07 times the twitch force in response to a singlet spike), indicating that doublets conferred a net advantage in force per spike only when twitch amplification was sufficiently strong (**Fig. 7h**, orange solid line). These simulations therefore delineate a regime in which doublets can enhance mean force efficiency without a proportional increase in spike output, providing a mechanistic basis for why such patterns may be selectively recruited under conditions of increased excitability. However, it comes at the cost of increased force variability per motor unit (**Fig. 7i, Extended Data Fig. 8**), with singlet firing varying considerably less. Since repetitive doublets only occur for motoneurons slightly above their threshold, they should represent a sub-pool of activated motoneurons, and this variability likely has little effect on total force output unless dense recruitment occurs at particular force levels.

Taken together, ultrafast ultrasound measurements showed that, for individual motor units, repetitive doublets are associated with twitch responses that exceed linear temporal summation in the two motor units examined, exhibiting gains greater than unity, where doublet firing can improve mean force without a proportional increase in spike output.

## Discussion

We identified a unified mechanism underlying motoneuron doublet firing. By integrating intracellular recordings, biophysically grounded simulations, and human motor unit data, we showed that both spikes of the doublet are initiated at the AIS, and that the second is driven by axonal NaP, which recurrently reactivates the AIS via currents returning from the axon that sum with dendritic and somatic Ca^2+^-mediated ADP, somatic NaP, and passive current. In the model, blocking any of these currents abolishes the doublet.

Early intracellular studies established Ca-mediated ADP as a key determinant of motoneuron doublet generation^21–23^. Classical interpretations attributed the ADP to the invasion of the action potential into electrotonically distant dendritic regions and the subsequent return of depolarizing current to the soma^30–32^. Our results refine this picture by clarifying the contributions of axonal and somatic NaP, Ca-mediated ADP and passive currents. Blocking any of these abolishes initial or repetitive doublets. Classical measurements indicate that the ADP time course is shaped in part by K conductance kinetics^66,67^, consistent with a fast-terminating K current that deactivates as the slower SK conductance engages and adds transiently to the depolarization. These K conductance kinetics would add to the depolarizing phase but would not remove the requirement for the axonal afterpotential. The requirement of an axonal afterpotential is supported by the literature on cortical neurons, where cutting proximal to the first node of Ranvier abolished doublets^37^. Although NaP and Ca conductances in dendrites are known to support bursting in cortical and hippocampal neurons^36,68,69^, our work suggests that dendrite- and axon-driven bursting is not due to two separate mechanisms^38^, but rather to one unified mechanism. However, our modeled dendritic compartment captures only the spatially averaged dendritic potential, which does not reach the local depolarization that activates Cav1.3 channels at discrete dendritic sites in real motoneurons, where dendritic Cav1.3 underlies the persistent inward current central to motoneuron excitability^70,71^. Importantly, our finding does not contradict this literature: in a spatially detailed dendrite, local Cav1.3 activation could serve the same post-spike timing function that the passive load provides in our reduced model, or contribute additional depolarizing drive. Resolving the relative contributions of passive load and active dendritic Cav1.3 will require a spatially extended model incorporating the dendritic tree.

For doublets, the requirement for axonal NaP is difficult to reconcile with pure dendritic-invasion models, which predict variable delays arising from electrotonic dispersion and state-dependent dendritic activation. Axons and nodes of Ranvier express NaP currents capable of sustaining depolarization and shaping excitability^72,73^. A passive contribution may add to this axonal afterpotential, since capacitive discharge of the internodal axolemma through the paranodal seal produces a depolarizing afterpotential that acts functionally like an inward nodal current^74–76^. This contribution would sum with the nodal NaP rather than replace it. This axonal afterpotential mechanism also provides a parsimonious link between repetitive doublet firing and classical F-wave physiology. Because F-waves reflect intrinsic motoneuron excitability following antidromic activation in the absence of synaptic input^77,78^, they place strong constraints on candidate mechanisms. A spike-activated axonal NaP current provides a sufficient mechanism by which an action potential can intermittently re-excite the motoneuron, potentially relating doublet firing during voluntary contractions to evoked F-wave responses. We found that at the threshold for evoking spikes (approximately - 55 mV), the ADP peak was below threshold (**Fig. 1d**, black line) and thus did not alone evoke a spike, explaining why F-waves are rare. In addition, the ADP with synaptic input was larger than that with antidromic stimulation (**Fig. 1f**), likely making doublets more common in natural contractions than after antidromic stimulation (F-wave). However, investigating this connection requires further research.

Earlier work proposed that initial and repetitive doublets arise from distinct mechanisms^10^. Our results instead support a unified mechanism in which a spike-activated axonal NaP current generates an ADP whose expression is governed by the balance of inward and outward currents relative to recruitment threshold. Thus, initial and repetitive doublets reflect different operating regimes of the same spike-activated axonal depolarizing process rather than distinct intrinsic mechanisms. This framework also prompts a reinterpretation of motor unit discharges during explosive ballistic contractions, such as fast force-and-hold tasks, in which the initial phase has long been characterized by instantaneous firing rates far exceeding those observed during sustained contractions^79^. These high rates have traditionally been interpreted as a transient increase in sustained discharge frequency driven by strong synaptic input. However, such instantaneous rates are indistinguishable from the doublet interval, and we propose that the apparent high-frequency firing at contraction onset in these tasks reflects the mechanism described here, including the passive properties (**Fig. 1h**, **Fig. 3l**).

Although both initial and repetitive doublets are prevalent, the shared mechanism we describe may nonetheless give them different roles in movement. Because our model indicates that a doublet follows when a synaptic input carries a motoneuron across threshold with a sufficient rate of rise, and rapid inputs are common in natural behavior, initial doublets may occur frequently during naturalistic contractions. Repetitive doublets, by contrast, require the Ca-mediated ADP together with the somatic and axonal NaP and passive currents to overcome the AHP on successive cycles, a balance that in healthy motoneurons is reached only near recruitment; in disease states that alter neuromodulatory drive or outward conductances, this balance may shift, and repetitive doublets could occur more broadly. These forms may also differ in their contribution to force. A doublet can in principle be produced by any unit that possesses the underlying machinery (the somatic and nodal persistent sodium currents and the Ca-mediated ADP) when it is driven rapidly across threshold, whereas repetitive doublets are, in the healthy state, largely confined to the small sub-pool near recruitment at a given moment. Motoneurons are likely to vary in this capacity since differences in NaP conductance or in axonal geometry, such as the position of the first node relative to the initial segment, would alter whether a doublet can be generated at all.

Despite the presence of a spike-activated axonal inward current, motoneurons rarely produce bursts beyond doublets in the repetitive-doublet regime. Our simulations indicate that NaP currents act as a transient, self-limiting amplifier of excitability. The first spike activates axonal NaP and generates an ADP sufficient to trigger a second spike, but the second spike occurs in a markedly altered biophysical context characterized by sodium channel inactivation, elevated intracellular calcium, and activation of potassium currents. This shifts the post-spike current balance toward stabilization, suppressing further delayed depolarizations and favoring doublets over triplets except possibly under conditions of strong neuromodulatory drive.

Once the repetitive doublets transitioned into single spiking, another motoneuron was typically recruited, producing either doublets or singlets (see **Fig. 6e**). This finding indicates continuous recruitment compensated by switching between singlet and doublet regimes, contrary to a more “stationary” model of motor unit behavior. One interpretation is that synaptic drive to the pool gradually increased to compensate for the declining output of adapting units, maintaining force while recruiting additional motoneurons as the intra-doublet frequency adapted and force-amplifying doublets were lost. However, several features argue against a substantial increase in common drive. The inter-doublet discharge rate of the repetitive-doublet units remained approximately constant across the contraction (**Fig. 6d-e**), in contrast to the doublet-frequency adaptation (**Fig. 6i**). Because the slower inter-doublet rate tracks the common drive more closely than the intrinsic intra-doublet interval, its stability suggests the drive changed little. The lower-threshold tonic units likewise showed stable or declining rates rather than the upward trend a rising drive would produce (**Fig. 6f-h**). Although adaptation could mask a modest increase in drive at the single-unit level, the stability of the less-confounded inter-doublet rate points to a relatively stable common drive. A stable drive cannot recruit new units by a simple threshold crossing; a potential explanation is that the effective recruitment threshold of the new unit declines over time, for example, through time-dependent facilitation (warm-up) of PICs under sustained monoaminergic drive. This would entail opposite processes acting in parallel: slow NaP inactivation driving intra-doublet adaptation in active units, and slow PIC facilitation lowering the threshold of unrecruited units, distinct conductances in different cells that need not be contradictory. Either way, the process underlying the doublet-to-singlet transition is likely slow, occurring over many seconds. Our data cannot definitively distinguish a small, adaptation-masked increase in drive from a stable drive with a falling recruitment threshold; this would require a direct estimate of the common drive across the plateau and an assessment of recruitment-derecruitment hysteresis. Regardless, the sequential recruitment and doublet-to-singlet transitions at constant force illustrate that steady output is maintained through dynamic reorganization of discharge across the pool rather than stable firing of individual units. The effect of repetitive doublets is likely negligible at most force levels since they occur only in a small fraction of units close to threshold. They may limit stability only during very low-force contractions. However, the effect of repetitive doublets on the force across force levels requires further investigation.

Our finding that the doublet-frequency adaptation is governed by the slow inactivation of the somatic NaP current suggests that this adaptation could serve as an experimentally accessible readout of an otherwise hard-to-measure intrinsic property. Because the adaptation time constant scales with the NaP inactivation time constant in our model, fitting the decay of the doublet-frequency adaptation may provide an indirect estimate of, or proxy for, the NaP inactivation kinetics in human motoneurons, which cannot be measured directly in vivo. This would offer a non-invasive window onto somatic NaP channel dynamics from surface or intramuscular recordings, with potential relevance for detecting altered excitability in neuromuscular disease.

Neuromodulation does not create the capacity for repetitive doublet firing but instead likely gates the expression of the somatic and axonal mechanism that is otherwise masked by outward conductances. By enhancing NaP currents and attenuating the AHP, neuromodulatory input can unmask these contributions to spike timing; in the model, this occurs within a parameter regime that preserves discrete firing rather than entering a plateau. This role complements, rather than replaces, the classical view of neuromodulation as amplifying synaptic gain through somatodendritic persistent inward currents^70,80^. To our knowledge, this specific mechanism, in which neuromodulatory facilitation of NaP shapes the temporal structure of motoneuron output through recurrent axonal contribution to the second spike, has not been described previously, suggesting an axis of neuromodulatory control that operates on spike timing in addition to firing rate and synaptic gain.

The functional relevance of doublet firing is underscored by its disproportionate effects on muscle force, traditionally attributed to enhanced Ca^2+^ release, accelerated cross-bridge recruitment, and increased short-range stiffness^14–16,81–84^. Our human motor unit recordings demonstrate that doublets can occur repeatedly during voluntary contractions and continue to produce nonlinear force enhancements, indicating that precise spike timing, in addition to average firing rate, plays a critical role in shaping motor output.

The presence of repetitive doublets in constant-force contractions also has implications for interpreting force steadiness. Force fluctuations during submaximal isometric contractions are classically attributed to low-frequency oscillations in the common synaptic input shared by the motoneuron pool, which dominate force variability^85,86^. At higher forces, most recruited units discharge well above threshold, where repetitive doublet expression is suppressed, and common input oscillations remain the principal determinant of variability. At very low forces, however, a substantial fraction of active motoneurons operate near the recruitment threshold, precisely the regime in which our results predict that repetitive doublets will emerge. In this regime, the spike-timing irregularity introduced by intermittent doublet firing would superimpose on common input oscillations and could constitute a previously unrecognized contributor to force variability, representing a potential trade-off of the same neuromodulatory drive that enhances force amplification. This interpretation may be particularly relevant during sustained constant-force contractions, where progressive recruitment of additional motor units is well documented and newly recruited units, discharging at low rates with unfused twitches, have been identified as a source of the increasing force variability observed as contractions approach task failure^87^. Because these units operate near their recruitment thresholds, our study suggests that they are especially likely to exhibit repetitive doublet firing, offering a complementary cellular explanation for the progressive loss of force steadiness during sustained contractions. Testing whether doublet prevalence tracks the decline in force steadiness across force levels and over the time course of sustained contractions represents a concrete prediction of our model.

Although long regarded as infrequent, doublet firing is likely heavily under-reported partly due to methodological biases. For example, multichannel surface EMG decomposition pipelines routinely exclude units with high instantaneous firing rates and use peak-detection methods that impose a minimum inter-spike interval on detected spike pairs, typically around 20 ms (50 Hz)^54,88^, which excludes the short intervals within a doublet. These peak-detection methods are amplitude-based and therefore prone to missing the smaller second spike of a doublet. The second spike of a doublet typically has reduced amplitude^64^, where the second peak of the decomposed spike train may be very close to the noise baseline (**Extended Data Fig. 9a-c**), increasing the likelihood of missed detections, particularly in decomposed spike trains with lower signal-to-noise ratios. Moreover, convolutive blind source separation algorithms are biased toward units with lower inter-spike interval variation. Whereas the coefficient of variation of the inter-spike interval averages about 15% for singlet-spiking units, it is far higher for units containing doublets, averaging over 100% and in some cases exceeding 200% (**Extended Data Fig. 9d-e**). When detection criteria are adjusted, doublets can be recovered^89^, and recordings during natural behavior show that initial doublets and high-frequency discharges are common types of firing during locomotion in both animals and humans^7,13^. Consistent with this, closely spaced spike pairs meeting the doublet criterion are clearly visible in published single-trial motor unit recordings from a macaque performing an isometric force task, although they were not examined as doublets in that study^56^. Initial and repetitive doublets appear in the slowly increasing and rapidly oscillating target forces, and during the fast sinusoids several units discharge recurring doublets, with both spikes of each pair assigned to the same unit by the original decomposition. This observation extends the phenomenon to voluntary force production in a non-human primate. Doublets in healthy humans have been observed using intramuscular EMG in a wide range of muscles, including tibialis anterior^11,13,64,90,91^, trapezius^8,12,92,93^, biceps^8,64^, triceps^8,93,94^, rectus femoris^12^, flexor carpi radialis^10^, flexor carpi ulnaris^24^, first dorsal interosseous^95^, soleus^96^, and the lumbar multifidus^97^. These observations, together with our findings, suggest that doublets are widely present in motoneuron firing and that their prevalence in human motor unit data from multichannel EMG over the last decades has been masked by methodological constraints. In contrast, some subjects may produce few or no repetitive doublets, perhaps due to dominant outward conductances or reduced neuromodulatory drive.

Some limitations should be considered when interpreting these findings. First, although our computational model captures key features of motoneuron excitability, it necessarily simplifies neuronal geometry; in particular, the lumped dendritic compartment cannot capture local dendritic Cav1.3 activation, and future targeted axonal recordings will be required to more precisely localize NaP conductances and determine how their spatial distribution interacts with dendritic and synaptic inputs. Second, while our intracellular recordings and pharmacological manipulations support an axonal origin of the ADP contribution underlying repetitive doublets, direct measurements of axonal NaP currents in motoneurons remain technically challenging and represent an important goal for future work. Third, we obtained the ultrasound-based observations of FRR in **Fig. 7** from a single subject due to substantial technical and experimental constraints. However, the remaining figures and the force-based simulations support these observations, and all converge on a coherent mechanistic interpretation linking doublet firing to nonlinear force enhancement.

In conclusion, we show that initial and repetitive doublets arise from the same mechanism. Both spikes of the doublet are initiated at the AIS. The second spike occurs when a slowly inactivating NaP current at the first node of Ranvier returns toward the initial segment and sums with dendritic and somatic Ca^2+^-mediated ADP, somatic NaP, and the passive membrane response, together overcoming the AHP. In the model, blocking any of these currents abolishes the doublet. This mechanism complements the classical role of neuromodulation in amplifying synaptic gain through somatodendritic persistent inward currents, revealing a further function in shaping the temporal structure of motoneuron output. More broadly, our results identify the axon as an active contributor to spike timing and intrinsic excitability, making it a dynamic target for motor control.

## Methods

### Animal experiments

Recordings were obtained from motoneurons and their axons in adult mice of both sexes in equal proportion, aged 2.5 to 6 months (see strain below). Mice were housed in groups of two to four under a 12 h light/dark cycle at controlled temperature and humidity, with unrestricted access to food and water. All procedures were approved by the University of Alberta Animal Care and Use Committee, Health Sciences division (ACUC protocols AUP00000224 and AUP00002891), and followed the guidelines of the Canadian Council on Animal Care. The ex vivo preparation and the recording, synaptic activation and stimulation procedures are summarized below and described in full previously^99,100^.

#### Ex vivo preparation of whole adult mouse sacral spinal cords

Mice were anaesthetized with urethane (0.11 g per 100 g body weight, to a maximum of 0.065 g). After laminectomy, the entire sacrocaudal spinal cord was excised rapidly and placed in oxygenated modified artificial cerebrospinal fluid (mACSF). All spinal roots were cut away apart from the sacral S3 and S4 and caudal Ca1 ventral and dorsal roots on both sides. The cord was held in the dissection chamber for 1.5 h at 20 °C and then transferred to a recording chamber perfused with normal ACSF (nACSF) at 23 to 32 °C and a flow rate above 3 ml/min. Residual anaesthetic was washed out over 1 h in nACSF before recording, after which the nACSF was recirculated in a closed system. The cord was pinned through connective tissue and cut root stumps onto a tissue paper bed at the base of a silicone elastomer (SilGuard) chamber. For recordings from motoneurons or V3 neurons, the cord was generally positioned dorsal surface up with the left side raised. This preparation is valuable because the small sacrocaudal cord is the only part of the adult spinal cord that remains viable as a whole ex vivo, allowing axonal conduction to be followed over long distances. The segment drives the axial musculature of the tail and provides a tractable model of motor function in intact and injured cords, with spinal circuitry, reflexes and motoneuron properties closely resembling those of hindlimb preparations^99,100^.

#### Electrode preparation and amplifier

Sharp intracellular electrodes were used so that motoneurons and axons could be recorded without injury or disturbance of their intracellular contents. Glass capillaries (603000, A-M Systems) were pulled on a Sutter P-87 puller (Flaming/Brown) and tip-filled with 2 M K-acetate combined with 2 M KCl in proportions giving KCl concentrations from 0 to 100 mM, or at 500 mM and 1000 mM, or alternatively with 500 mM KCl in 0.1 M Trizma buffer containing 5 to 10% neurobiotin (Vector Laboratories). Tips were then beveled to 30 MΩ on a rotary beveler (BV-10, Sutter Instrument).

Electrodes were advanced into motoneurons in 2 µm steps while brief high-frequency current pulses (capacitance overcompensation) and audio feedback guided penetration. Cells were identified as motoneurons from the antidromic response to ventral root stimulation, together with ventral horn position, input resistance and a membrane time constant above 6 ms.

Recording and current injection used an Axoclamp-2B amplifier (Axon Instruments/Molecular Devices). Signals were low-pass filtered at 10 kHz and digitized at 30 kHz (Clampex and Clampfit; Molecular Devices). For some measurements, the amplifier was run in discontinuous single-electrode voltage-clamp mode (gain 0.8 to 2.5 nA/mV; for Ca PICs) or, during current injection, in discontinuous current-clamp mode (switching rate 7 kHz).

Axons were usually impaled in the myelinated internodal region rather than at a node, since nodes occupy only a small fraction of the axon and are correspondingly unlikely to be penetrated. When threshold was tested with current pulses (20 ms, rheobase test), the spikes of the two nodes flanking the electrode could be resolved separately, because near threshold the pulse sometimes evoked a spike at only one node, roughly halving the total spike amplitude, as expected if the electrode lay about midway between two nodes separated by roughly one space constant.

#### Activating motoneurons synaptically with V3 activation

Motoneurons were driven synaptically by optogenetic stimulation of V3 neurons in a strain of mice with Cre recombinase expressed under the Sim1 promoter region, since the definition of V3 neurons is expression of Sim1 during development. The optogenetic stimulation of the Sim1-Cre mice (Sim1-Cre-ki and Sim1-Cre-tg)^99^ was performed using 447 nm (D442001FX) and 532 nm (LRS-0532-GFM-00200-01) lasers (Laserglow Technologies, Toronto), as described in detail previously^99^. Laser light passed through a fiber optic cable to a half-cylindrical prism spanning about two spinal segments (8 mm; focal length 3.9 mm, Thorlabs, Newton), which shaped it into a narrow beam 200 µm wide and 8 mm long. The beam was usually aligned longitudinally along the left side of the cord to recruit many V3 neurons at once. Because Cre-driven channelrhodopsin-2 (ChR2) depolarizes neurons rapidly, 5 to 10 ms light pulses were sufficient to activate them. Intensity was set to three times threshold (3xT), where threshold is the light level needed to evoke a response and 3xT approaches the maximal response; this level penetrated the full thickness of the cord while remaining low enough to avoid local heating. In control tests, monosynaptic activation of motoneurons, mediated by the most ventral V3 neurons that synapse directly onto them, appeared at 2.3xT, confirming that 3xT drove the entire width of the cord, though with reduced intensity at depth. Functional ChR2 in motoneurons was confirmed directly by advancing electrodes into cells that responded to light (5 to 10 ms pulse, 447 nm) with no synaptic delay (under 1 ms) and a response that persisted after synaptic transmission was blocked.

#### Ventral root stimulation

Ventral roots were laid on silver-silver chloride wires positioned just above the nACSF and sealed under grease (petroleum jelly and mineral oil, 3:1) for monopolar stimulation. A surrounding layer of more viscous high-vacuum grease prevented oil from leaking into the bath flow. Bipolar stimulation was used at times to reduce the stimulus artifact when recording from ventral roots. Roots were driven by a constant current stimulator (ISO-Flex) with 0.1 ms pulses.

#### Ventral root grease-gap recording

A grease-gap method was used to capture the summed intracellular response of many axons through ventral root recordings. Freshly cut roots were placed on silver-silver chloride wires just above the bath and sealed in grease over 2 to 5 mm, with longer coverage giving a tighter seal; grease was applied as close to the cord as possible to maximize the signal. Reference and ground wires, also silver-silver chloride, were placed in the bath.

Composite EPSPs across many motoneurons were similarly recorded from the central cut ends of the S3 and S4 ventral roots sealed in grease. EPSPs were identified as monosynaptic from their rapid onset, low-latency jitter (under 1 ms), persistence at 10 Hz stimulation, and isolated appearance at the threshold for evoking EPSPs by ventral root stimulation. EPSPs were usually elicited by stimulating the Ca1 or S4 roots. Recordings were amplified 2000 times, high-pass filtered at 0.1 Hz to remove drift, low-pass filtered at 10 kHz and digitized at 30 kHz (AxoScope 8; Axon Instruments/Molecular Devices).

#### Drugs and solutions

Two artificial cerebrospinal fluids were used: mACSF in the dissection chamber before recording and nACSF in the recording chamber. mACSF contained (in mM) 118 NaCl, 24 NaHCO_3_, 1.5 CaCl_2_, 3 KCl, 5 MgCl_2_, 1.4 NaH_2_PO_4_, 1.3 MgSO_4_, 25 _D_-glucose and 1 kynurenic acid. nACSF contained (in mM) 122 NaCl, 24 NaHCO_3_, 2.5 CaCl_2_, 3 KCl, 1 MgCl_2_ and 12 _D_-glucose. Both were saturated with 95% O_2_ and 5% CO_2_ and held at pH 7.4. Drugs added to the ACSF in some experiments included 5-HT (Sigma-Aldrich) and TTX (TTX-citrate; Toronto Research Chemicals, Toronto). Stocks were prepared at 10 to 50 mM in water or DMSO and diluted to the final concentration in ACSF.

### Three-compartment conductance-based motoneuron model

To investigate the ionic mechanisms underlying doublet discharges in human motoneurons, we implemented a three-compartment, conductance-based model comprising a dendrite, soma, and axon. The model is based on Hodgkin-Huxley formalism and was adapted from the single-compartment motoneuron model^47^, which was originally constrained by patch-clamp recordings from neonatal rat hypoglossal motoneurons. A subset of maximal conductances was re-scaled to yield firing rates consistent with those reported for human motoneurons recorded in vivo, while preserving the biophysical structure of the original model. The three compartments are electrotonically coupled through symmetric axial conductances, with the soma positioned between the dendritic and axonal compartments. Each pair of neighboring compartments was coupled by a single conductance g_c_, such that the current flowing from compartment X into compartment Y is I = g_c_(V_X_-V_Y_); the dendrite-soma and soma-axon couplings (g_c,ds_ and g_c,sa_) were specified independently. Stimulus current was injected into the somatic compartment.

#### Ionic currents

The somatic compartment contains the full complement of ionic currents of the original single-compartment model^47^ that inspired the present model: a fast transient sodium current (I_NaF,s_), a NaP current (I_NaP,s_), a delayed-rectifier potassium current (I_K,s_), a transient A-type potassium current (I_A,s_), an SK current (I_SK,s_), a hyperpolarization-activated cation current (I_H,s_), three voltage-gated calcium currents of the P-, N-, and T-type (I_P,s_, I_N,s_, I_T,s_), and a passive leak current (I_leak,s_). Note that we refer to the P- and N-type Ca currents as the HVA Ca currents and the T-type current as the LVA Ca current. Intracellular calcium followed a first-order buffering equation that relates total calcium current to I_SK,s_ gating. We lumped the somatic compartment with the AIS.

The axonal compartment contains currents tailored to spike propagation and regeneration: a fast axonal sodium current (I_NaF,ax_), a delayed-rectifier potassium current (I_K,ax_), a NaP current (I_NaP,ax_), and a passive leak (I_leak,ax_)^101^. This compartment represents the first node of Ranvier. The axonal NaP current, which we identify as necessary for doublets, is described by the same voltage-dependent activation form as its somatic counterpart but with gating kinetics adjusted to the axonal regime (inactivation time being 150 ms in the soma and 25 ms in the axon, see Supplementary Methods).

The dendritic compartment was treated as an active, non-spiking compartment. It contained a dendritic SK current (I_SK,d_) gated by a separate dendritic calcium pool, dendritic N- and T-type calcium currents (I_N,d_, I_T,d_) with their own gating variables, and a passive leak (I_leak,d_). Fast sodium, delayed-rectifier, and A-type potassium currents are absent from the dendrite, in agreement with the low density of action-potential-generating conductances observed in motoneuron dendrites. Dendritic calcium dynamics was governed by its own buffering equation, independent of the somatic calcium pool, so that dendritic SK activation was driven only by local dendritic calcium entry. We note that, because the dendrite was represented as a single lumped compartment, its spatially averaged membrane potential does not reach the local depolarization required to substantially activate the higher-threshold dendritic calcium currents; the consequences of this simplification for the dendritic contribution to doublets are addressed in the Discussion.

All gating variables followed a first-order kinetic scheme, with steady-state activation/inactivation functions and time constants taken from a previous single-compartment model^47^. The complete set of kinetic expressions, reversal potentials, maximal conductances, and initial conditions is provided in the Supplementary Methods.

#### Numerical integration and stimulation protocols

The coupled ordinary differential equations of the three-compartment model were integrated using a fixed-step fourth-order Runge-Kutta (RK4) scheme implemented in MATLAB (MathWorks, Natick, MA, USA), with an integration step of Δt = 1 µs. At each step, gating variables were constrained to [0,1], and intracellular calcium concentrations were constrained to be non-negative to avoid numerical drift in long simulations.

To mimic stimulation protocols routinely used in whole-cell patch-clamp recordings from mammalian motoneurons, the injected current was defined as a two-step waveform: a constant holding current applied for a long pre-stimulus interval, followed by a rectangular step current of variable amplitude applied to the somatic compartment. The pre-stimulus interval was chosen to be sufficiently long (2.5 s) for all state variables to reach steady state under the holding current, so that the neuron’s response to the step current was independent of the numerical initial conditions. Then, the first 1.5 s was discarded. The step duration was varied across experiments: a 1 s step was used for the standard firing pattern and blocking analyses, and a 10 s step was used to quantify doublet-frequency adaptation.

A complete list of parameters, together with their nominal values and the ranges explored, is provided in the Supplementary Methods.

### Human experiments

Two human experiments were conducted involving the lower leg and the tibialis anterior muscle with dorsiflexion tasks. The first experiment included one HDsEMG grid with a steady-force task comprising trapezoidal paths and a fast-contraction task comprising sinusoidal paths. The second experiment included HDsEMG and ultrafast ultrasound, with only steady-force tasks, enabling quantification of the neuromechanics associated with motoneuron spiking.

#### Subjects

In experiment 1, fourteen healthy subjects were included in the study (8 males, 30 ± 7 years; 6 females, 30 ± 5 years). In experiment 2, twelve healthy subjects were included in the study, although ten subjects (26.2 ± 2.9 years) were used for the analysis due to system and synchronization errors. We numbered the subjects from #1 to #24, with the first 14 subjects from experiment 1, and the additional 10 subjects from experiment 2. The experimental protocol was clearly explained to the subjects before each experiment, and they provided informed consent, both orally and in writing. These experiments were approved by the Swedish Ethical Review Authority (experiment 1, reference: 2025-01204-01) and Imperial College Research Ethics Committee (experiment 2, reference: 20IC6422) in accordance with the Declaration of Helsinki.

#### Instrumentation

The HDsEMG signals were recorded using 64-electrode grids (13x5 formation with 8 mm inter-electrode distance, one electrode in the corner removed; OT Bioelettronica, Torino, Italy), where the EMG signals were recorded in monopolar derivation, amplified, sampled at 2000 Hz (experiment 1) or 2048 Hz (experiment 2), A/D converted to 16 bits with 150x gain, and bandpass filtered between 10 Hz and 500 Hz using a Sessantaquattro+ (experiment 1) or Quattrocento (experiment 2) amplifier (OT Bioelettronica, Torino, Italy). The forces produced by dorsiflexion were recorded using an ankle dynamometer and transmitted via a Forza force amplifier (OT Bioelettronica, Torino, Italy) to the respective amplifier.

The ultrasound signals were recorded using the Vantage Research Ultrasound Platform (Verasonics Vantage 256, Kirkland, WA, USA) through an L11-4v transducer with 128 elements and a center frequency of 7.24 MHz. The transmit-receive protocol employed single-angle plane-wave imaging at 1 kHz for 30-second recordings. To synchronize the ultrasound and EMG systems, the Verasonics system produced a one-microsecond pulse, which was elongated by an Arduino Uno and fed into the Quattrocento Amplifier to allow for alignment offline.

#### Experimental procedures

The skin area above the tibialis anterior muscle was shaved when necessary and cleansed with an abrasive paste. The HDsEMG electrode grid in experiment 1 was attached to the skin after palpation. In experiment 2, two electrode grids were used, with the ultrasound probe positioned between them. The first electrode grid was placed proximally, parallel to the muscle fibers, whereas the second electrode grid was placed distally, leaving a gap of approximately 1 cm between the grids. The electrodes were secured with tape and, where necessary, self-adhesive medical bandages. The ultrasound probe was placed in a custom-designed 3D-printed probe holder and attached to the leg in the gap between the EMG grids with the imaging plane perpendicular to the muscle fibers. As such, the resultant images were cross-sectional views of the tibialis anterior muscle. A water-based ultrasound gel improved the coupling between the probe and the leg.

The subject was seated at a comfortable distance from a computer screen, and their leg was secured in an ankle dynamometer with the foot at a 90-degree angle to the leg. Soft padding was placed around the leg to provide comfort and to secure it in position.

Real-time force feedback from the ankle dynamometer was provided to guide the subject along on-screen force paths. Initially, the subject was instructed to perform up to three of their strongest dorsiflexion contractions, and the highest force was recorded as their MVC. Thereafter, they performed different tasks. In experiment 1, they performed five 60-second ramp contractions (5, 10, 15, 20, and 25% MVC) shaped like a trapezoid, with the first and last 5 s at 0% MVC, a 5 s ramp up, a 40 s plateau, a 5 s ramp down, and 5 s at 0% MVC. After the steady trapezoidal contractions, they performed two fast sinusoidal contractions at a 2 Hz rate (a ramp up to 10% MVC and superimposed sinusoids between 5-15% MVC). Some subjects also performed 1 Hz contractions, as shown in **Fig. 4**, and some subjects also performed triangular contractions (0 to 30% MVC over a 15-second ramp). For the doublet occurrence, we analyzed only one of the two recordings. For all subjects except one, this was the second recording, as the first was used to acclimate to the task.

In experiment 2, they performed four ramp contractions at 2%, 5%, 10%, and 20% MVC, similar to experiment 1 with 5-second ramps and a 40-second plateau. Once the subject reached the plateau and the force was judged stable, the experimenter began a 30-s ultrasound recording. Between tasks, the subject rested for up to 60 s to ensure no fatigue.

#### HDsEMG data processing

The EMG signals in each recording were band-pass filtered from 20 to 500 Hz using a zero-phase 2nd-order Butterworth filter, along with a 50 Hz notch filter, before being fed into the decomposition method^54^ within the open-source MUedit software^88^. The recordings were decomposed sequentially, i.e., starting with the smallest force levels (5% in experiment 1 and 2% in experiment 2), moving on to higher force levels. The estimated spike trains were manually curated according to published guidelines^88,102^. Once manually curated, we extracted the motor unit action potentials (with a -25 to 25 ms window) and used them to construct a motor unit filter^89,103^ that was applied to the next recording (10% MVC in experiment 1 and 5% MVC in experiment 2), where the extracted sources from this constructed motor unit filter were stored along with the already estimated sources from the algorithmic decomposition. These were, in turn, manually curated before constructing motor unit filters and applying them to the next recording, and so on. This procedure was performed to maximize the motor unit yield.

To estimate the number of uniquely identified motor units for each subject across all recordings, we computed cross-correlations between the stored, edited, clean sources and the motor unit filter-based sources extracted from them. If the cross-correlation was >0.65, we considered it a match. This cutoff was determined through empirical testing in different recordings, by comparing firing rates and spatiotemporal action potential shapes between different recordings from the same subject. This procedure yielded unique motor unit action potentials for each subject, and from these, we could repeat the motor unit filter approach to extract sources from the fast sinusoidal contractions.

In the manual curation process, we reduced the minimum distance between peaks to a value smaller than that used by the standard MUedit implementation. As the manual curation process is subjective in nature, the criteria for a doublet were set to be within 10 ms (100 Hz or higher) between two spikes, and the inter-spike interval should be prolonged compared to between two single spikes^64^.

The recruitment threshold (in terms of % MVC) was computed as the MVC level at which the mean of the first three inter-spike intervals fell below 250 ms.

#### Ultrasound data processing

The raw radiofrequency signals were beamformed using the delay-and-sum method, yielding 30,000 frames of 357x128 pixels (depth vs. lateral, corresponding to approximately 40x38.4 mm). To quantify intramuscular displacements (in terms of velocity) in response to neural discharges, phase shifts within 1-mm depth windows were computed for radiofrequency signals between two consecutive frames^59^.

To compute the region of the muscle cross-section that produced synchronous displacements in response to neural discharges, we used spike-triggered averaging (STA), triggering the velocity image sequences with the obtained motoneuron spike train^62^. This procedure resulted in 100-millisecond spatiotemporal sequences (50 milliseconds before and after the trigger). To penalize large variabilities in velocity across triggers, the squared STA was computed, and the STA variance was divided by the variance of the curves comprising the STA, for each pixel in the image series. Thus, regions with high velocities and low variability have high STA magnitudes. The STA magnitudes were modulated with a negative sign if the direction of the motion was away from the probe at the time of the spike. Next, we selected the 10 pixels with the largest STA magnitude (referred to as the ROI in **Fig. 7**) and calculated the average twitch profiles. In this study, positive velocity indicated displacement toward the probe and skin, whereas negative velocity indicated displacement in the opposite direction.

#### Statistical analysis

Doublet-frequency adaptation during repetitive doublets was quantified by fitting an exponential decay model to instantaneous firing rates (see **Fig. 5g**, **Fig. 6i**). For each motoneuron, the firing rates were modeled as *f*(*t*) = *ae^bt^* + *c* by minimizing the sum of squared residuals with the Nelder-Mead simplex algorithm (MATLAB fminsearch). See Section S3 in Supplementary Methods for more details. For the human experimental data, we retained only fits that converged for further analysis. To enable comparison across motor units and subjects, fitted profiles were aligned to the onset of the first doublet (t = 0) and normalized to the initial firing rate. Population-level adaptation curves were obtained by averaging normalized profiles over time.

Statistical equivalence to 1 was assessed using a nonparametric bootstrap implementation of the two one-sided tests (TOST) procedure (see **Fig. 6k**). The median was used, and its sampling distribution was estimated from 10,000 bootstrap resamples with replacement. Equivalence bounds were defined a priori as ±5% ([0.95, 1.05]). Statistical equivalence was concluded if the bootstrap (1-2α) confidence interval (90% for α = 0.05) lay entirely within the equivalence region.

Paired differences in firing rate between singlet and repetitive doublet spiking were assessed using a Wilcoxon signed-rank test, a nonparametric test for median differences (see **Fig. 6l**). For each motor unit, the difference between singlet and doublet firing rates was computed, and the null hypothesis of zero median difference was tested. Statistical significance was assessed at α = 0.05.

#### Twitch force modeling

To quantify the effect of doublet firing on force production, we simulated unfused tetanic contractions of a single motor unit using stochastic spike trains and a physiologically based twitch model^65^. Spike trains were generated over 10 s at a sampling rate of 1 kHz, with inter-spike intervals drawn from a normal distribution (coefficient of variation = 10%). For each stochastic realization, singlet firing rates were sampled from a uniform distribution over 11.2 to 13.6 Hz, matching the physiological range of singlet discharge rates measured in our human motor unit recordings, and the corresponding doublet firing rate was set to a random proportion (56 to 75%) of the singlet rate, reflecting the observed ratio of doublet to singlet discharge rates in the same recordings, and matched observations in the literature^64^. Twitch contraction time was sampled from a uniform distribution over 75 to 90 ms^105^, and the half-relaxation time was set to 5/3 of the contraction time^106^, consistent with reported motor unit twitch properties. Doublets were modeled by inserting a second spike 5 ms (corresponding to 200 Hz, which is approximately what we observed in our human data, see **Fig. 6e**) after each primary discharge, matching the intra-doublet interval observed experimentally. Force output was computed by convolving spike trains with the twitch response, and the amplitude of twitch responses during doublets was scaled by a gain factor ranging from 0.5 to 1.5 in steps of 0.01 to simulate amplification of force. For each gain value, 100 stochastic realizations were generated. Mean force and force normalized to the number of spikes were computed for singlet and doublet conditions, and differences between conditions were averaged across realizations (see **Fig. 7h**).

## Acknowledgements

We would like to thank Erik Stålberg and Maria Piotrkiewicz for discussions prior to this paper, which ultimately guided us in this direction.

## Author contributions

Conceptualization (R.R., D.B., M.G., D.F.), Methodology (R.R., D.F.), Software (R.R.), Validation (R.R.), Formal analysis (R.R.), Investigation (R.R., D.B.), Resources (R.R., D.F.), Data Curation (R.R., D.B.), Writing – Original Draft (R.R.), Writing – Review & Editing (R.R., D.B., M.G., D.F.), Visualization (R.R., D.B.), Supervision (D.F.), Project administration (R.R.), Funding acquisition (R.R., D.F.).

## Code and Data Availability

The data supporting the findings of this study are available on Harvard Dataverse at https://doi.org/10.7910/DVN/9YZROY, under a CC BY 4.0 license. The code used to generate the figures, including the three-compartment simulation model, is available on GitHub at https://github.com/robinrohlen/doublet-paper, under an MIT license.

## Extended Data Figures

**Extended Data Figure 1.**
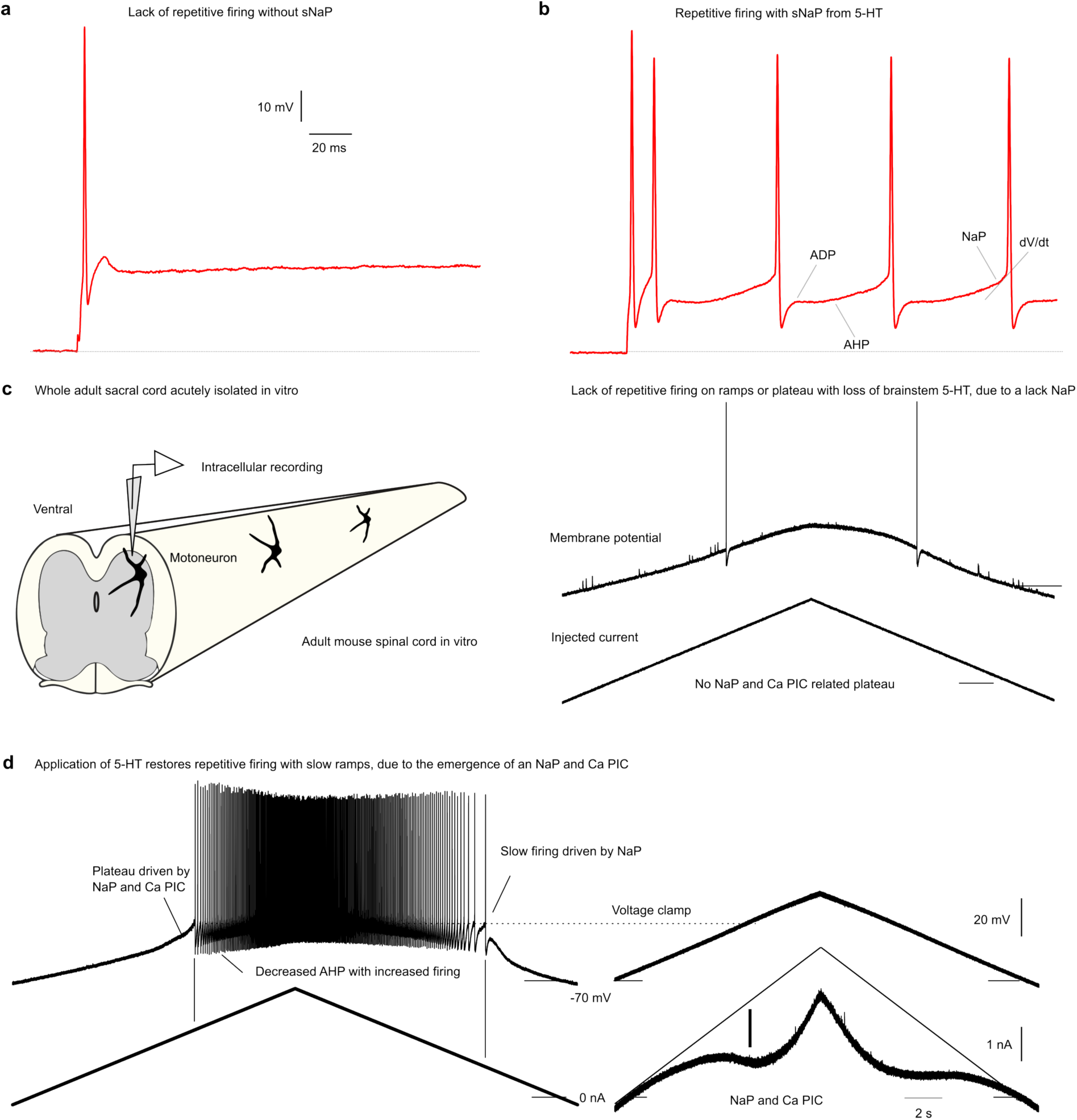
| Somatic NaP is crucial for repetitive firing. **a** and **b,** Fast-rising steps could initiate single spikes but generally could not initiate repetitive firing unless the cell possessed a soma NaP. **c**, Likewise, during ramp current injections, repetitive firing generally required a soma NaP, as evidenced by the lack of NaP (a linear I-V response) and of repetitive firing in the acutely isolated spinal cord, where the brainstem monoamines required for the current are removed. **d**, In contrast, facilitating the NaP by adding 5-HT to replace lost brainstem drive enabled repetitive firing, even at long intervals where the AHP had ended (slow firing). This NaP was quantified during voltage clamp, where it was initiated subthreshold to spiking and, together with the slower-rising Ca^2+^ persistent inward current (Ca PIC), produced a plateau potential that accelerated the membrane toward spike threshold and thereby initiated firing.

**Extended Data Figure 2.**
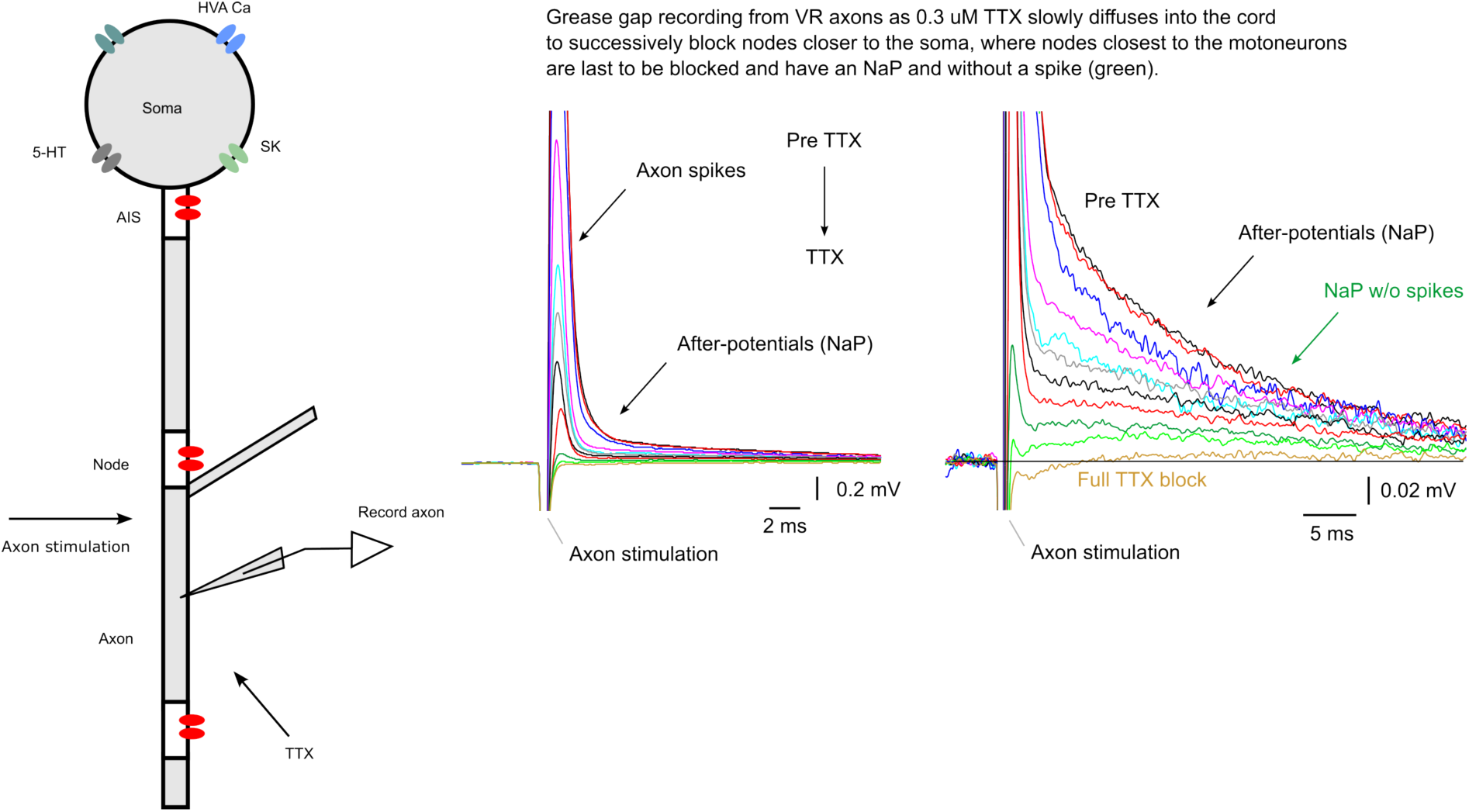
| Axon spike shape and the NaP origin of its long tail current. A long afterpotential was evident when recording from multiple axons in the ventral root using the grease-gap method, while the axons were stimulated at the ventral root entry zone. Application of low-dose TTX (0.3 µM) slowly blocked the NaP and spikes, progressively reducing the composite action of the afterpotential in discrete steps as TTX diffused to nodes deeper into the spinal cord. Some axons exhibited an afterpotential in the absence of a spike, and this was eventually blocked by TTX (green traces).

**Extended Data Figure 3.**
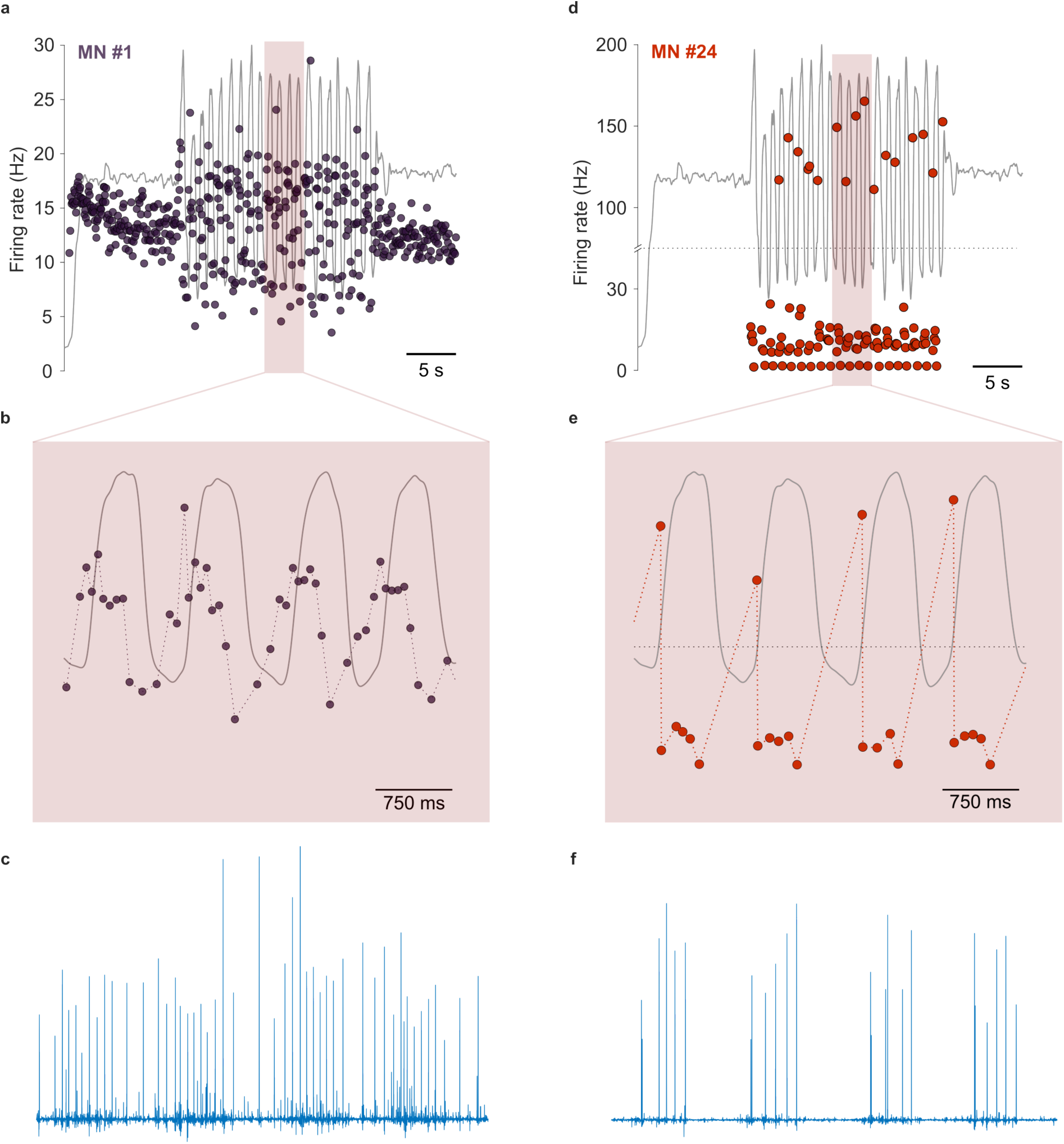
| Firing patterns and separated spike train sources from humans during fast sinusoidal contractions. **a** and **b**, The firing pattern from motoneuron (MN) #1 in Fig. 4 exhibited phase advance relative to the force oscillations. **c**, The separated spike train source in (**a**-**b**) from the EMG signals shows a high signal-to-noise ratio. **d** and **e**, The firing pattern of MN #24 in Fig. 4 of the main paper produced a doublet preceding each rapid increase in force. **f**, The separated spike train source in (**d**-**e**) from the EMG signals shows a high signal-to-noise ratio.

**Extended Data Figure 4.**
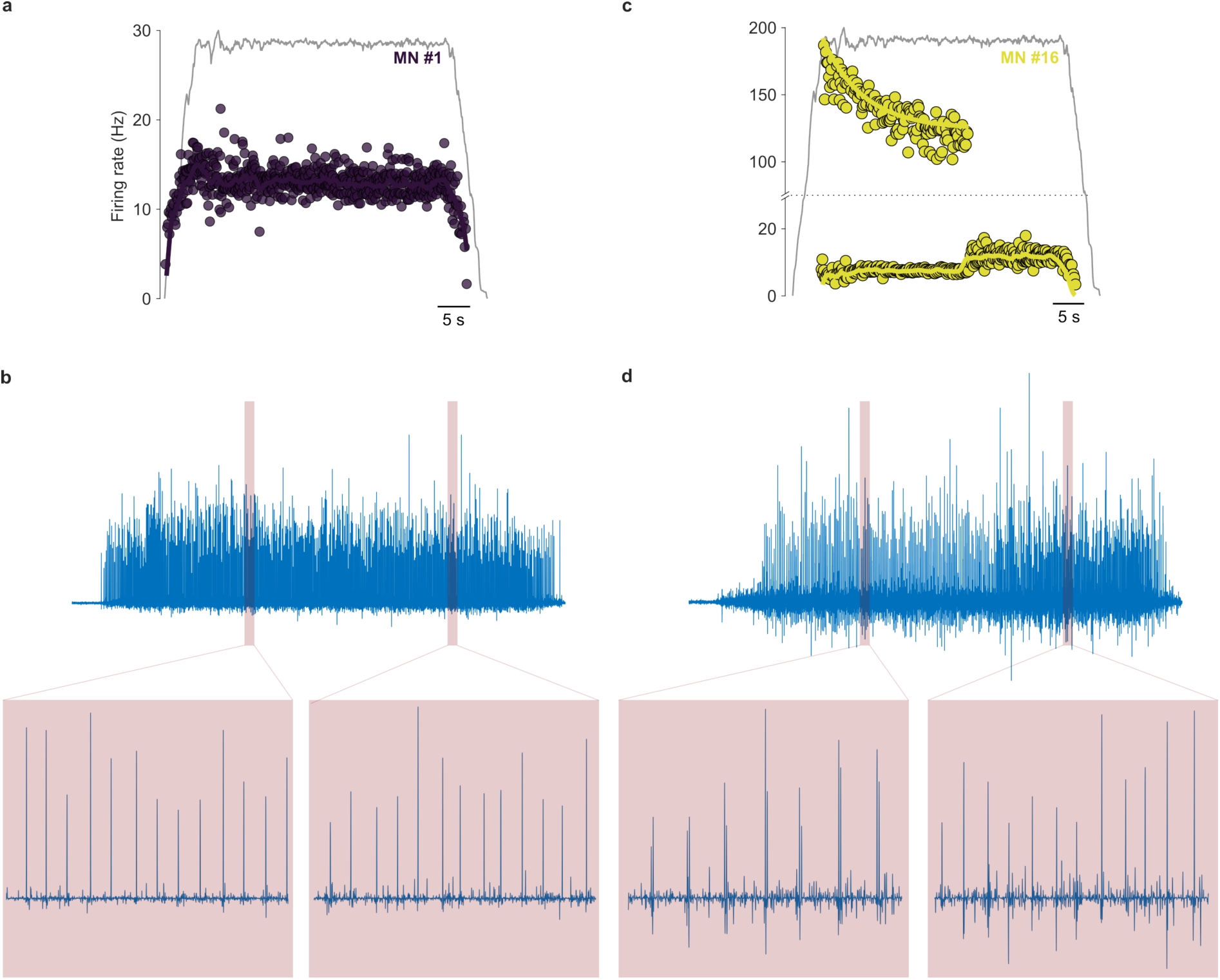
| Firing patterns and separated spike train sources from humans during stable contractions. **a**, Representative firing profile from subject #13 in Fig. 6: a force-correlated spike train. **b**, The separated spike train source in (**a**) from the EMG signals shows a high signal-to-noise ratio. **c**, Another representative firing profile: repetitive doublets that transition into singlet spiking. **d**, The separated spike train source in (**c**) from the EMG signals shows a high signal-to-noise ratio for both the doublet spikes and singlet spikes.

**Extended Data Figure 5.**
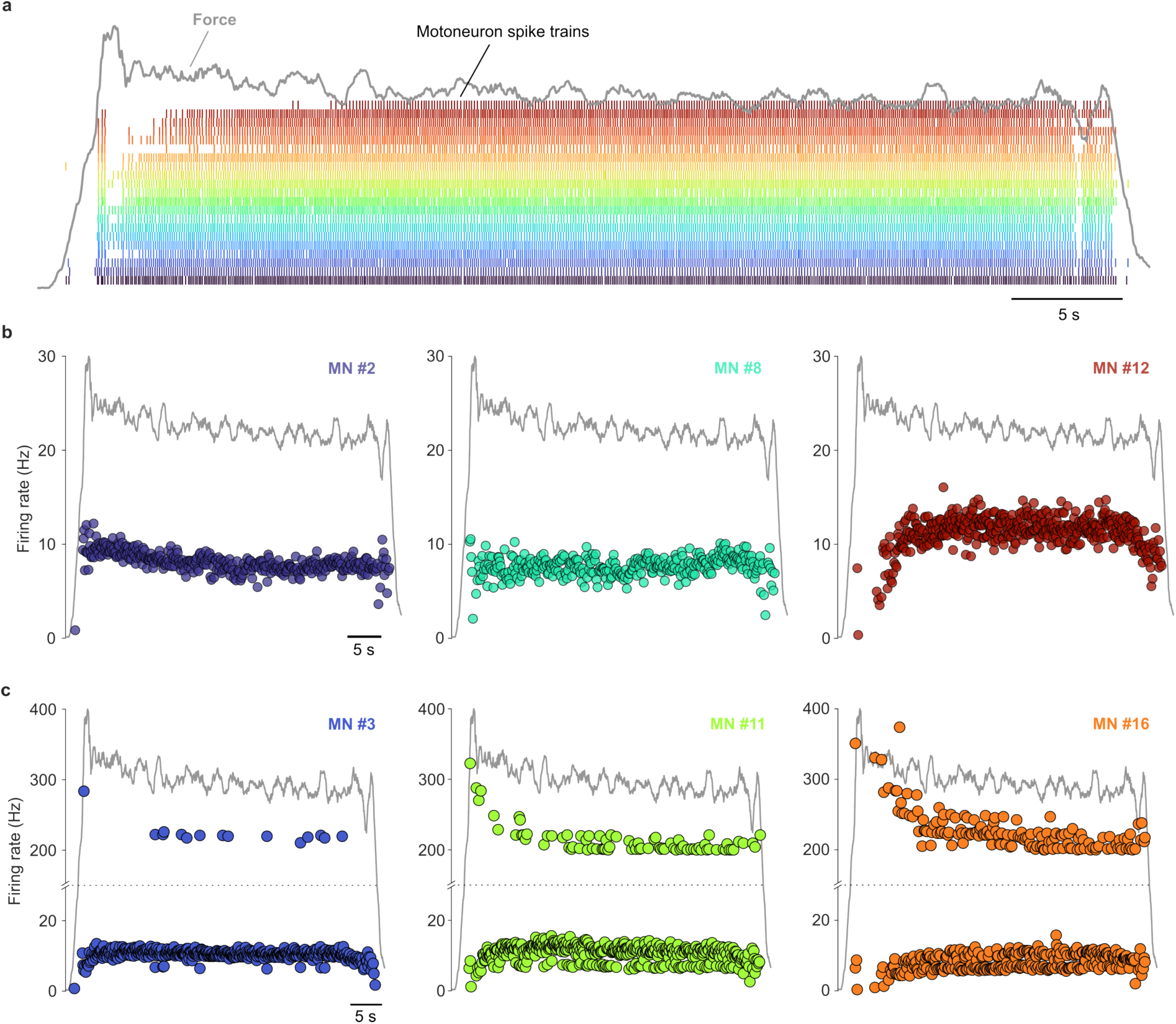
| Human doublets intermingled throughout a stable isometric contraction. **a**, The identified spike trains in subject #2 performing a trapezoidal isometric contraction in a dorsiflexion task at 5% MVC. **b**, Singlet firing patterns of three motoneurons. **c**, Doublet firing patterns showing singlet and doublet spikes intermingled throughout the contraction.

**Extended Data Figure 6.**
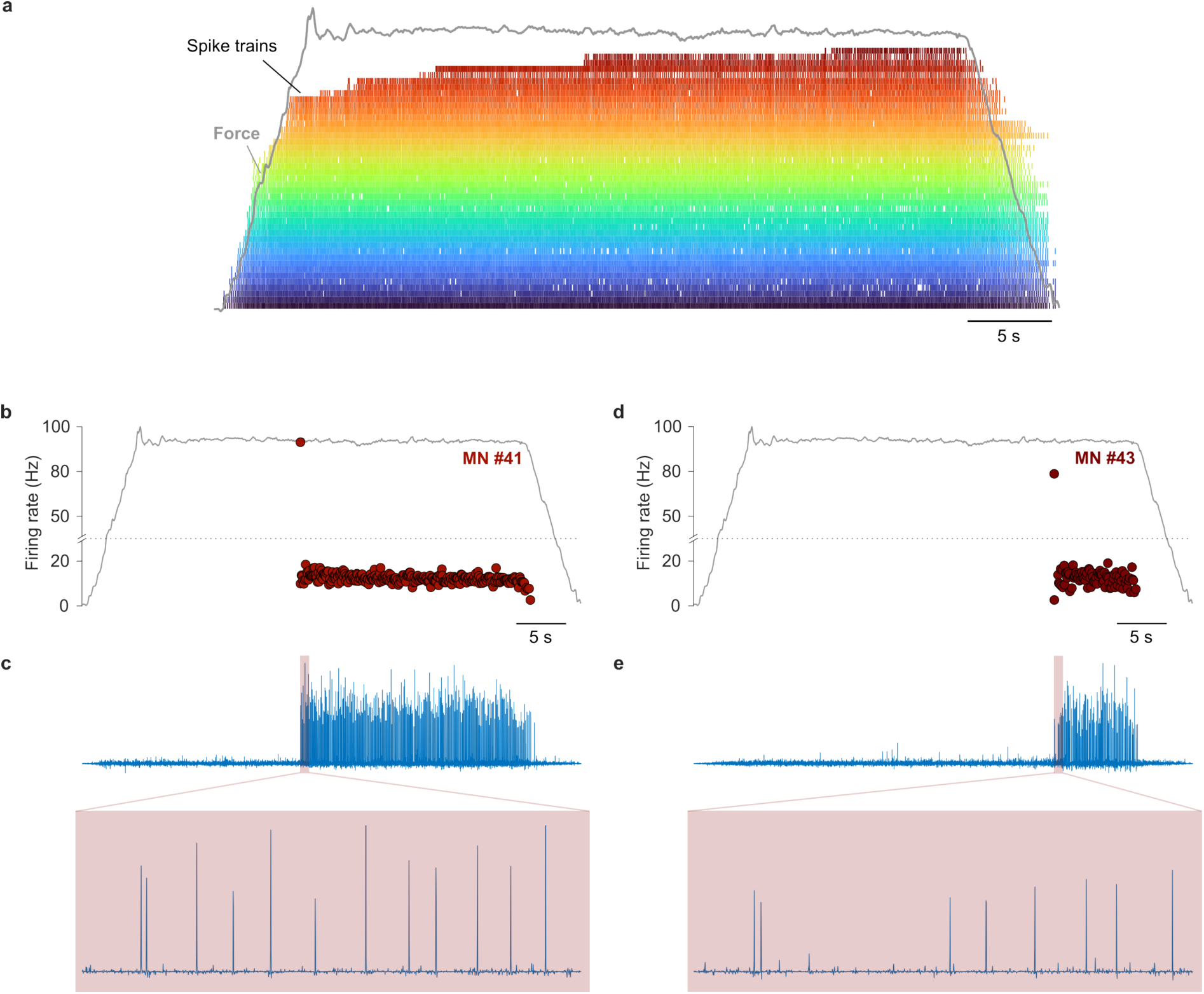
| Human doublets with an instantaneous firing rate below 100 Hz. **a**, The identified spike trains in subject #1 performing a trapezoidal isometric contraction in a dorsiflexion task at 25% MVC. **b**, Firing pattern of MN #41 with an initial doublet having an intra-doublet interval of 11 ms (instantaneous frequency of 91 Hz). **c**, The separated spike train source in (**b**) from the EMG signals showing a prolonged inter-spike interval. **d**, Firing pattern of MN #43 with an initial doublet having an intra-doublet interval of 13.5 ms (instantaneous frequency of 74 Hz). **e**, The separated spike train source in (**d**) from the EMG signals showing a prolonged inter-spike interval.

**Extended Data Figure 7.**
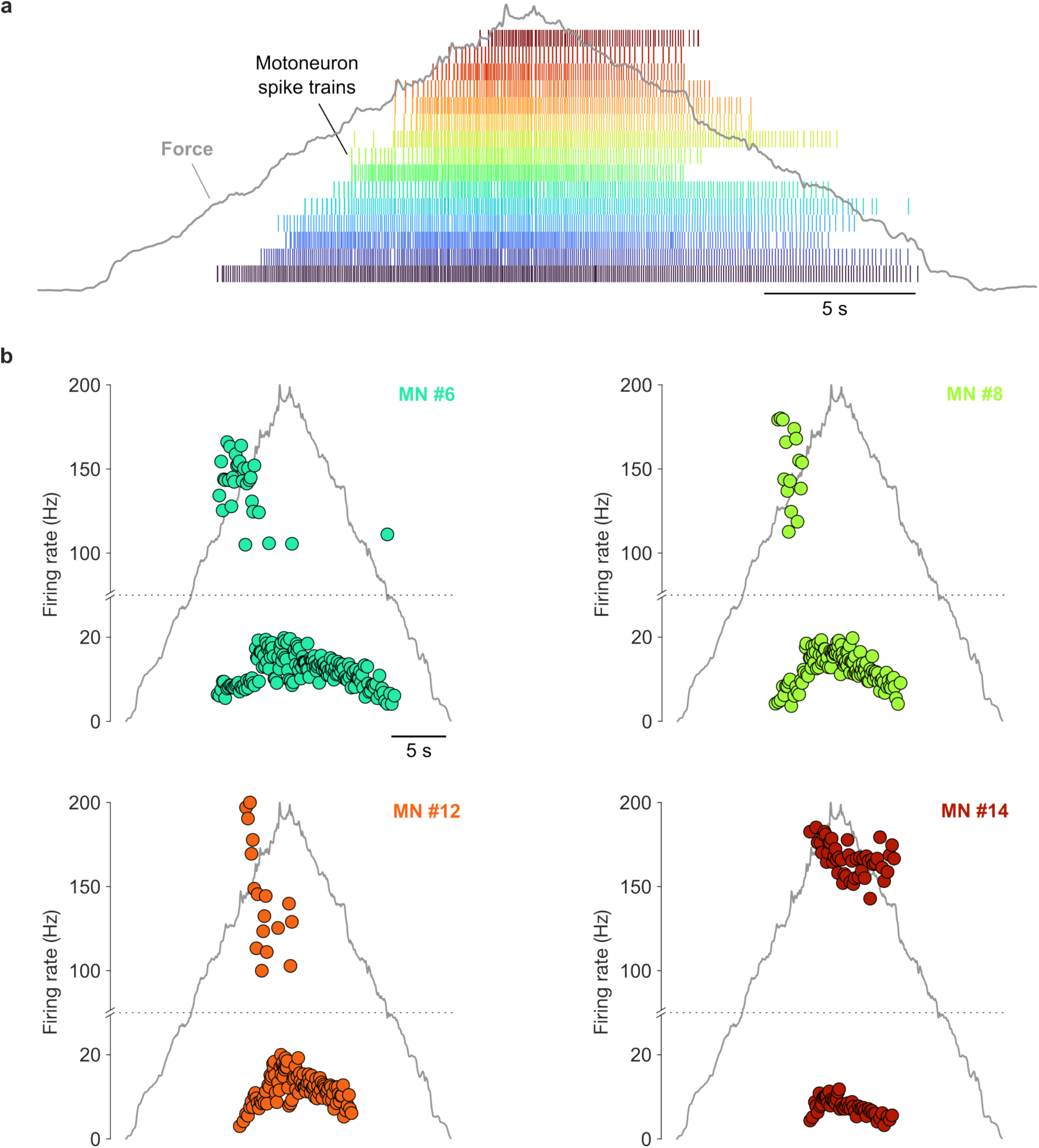
| Human doublets during a triangular contraction. **a**, The identified spike trains in subject #13 performing a triangular isometric contraction in a dorsiflexion task from 0 to 30% MVC with 15-second ramps. **b**, Doublet firing patterns for motoneuron (MN) #6, #8, #12, and #14. Note that the doublets typically occur within a close force level and show an increased intra-doublet interval over time. MN #14 only produced doublets and was recruited close to the 30% MVC level.

**Extended Data Figure 8.**
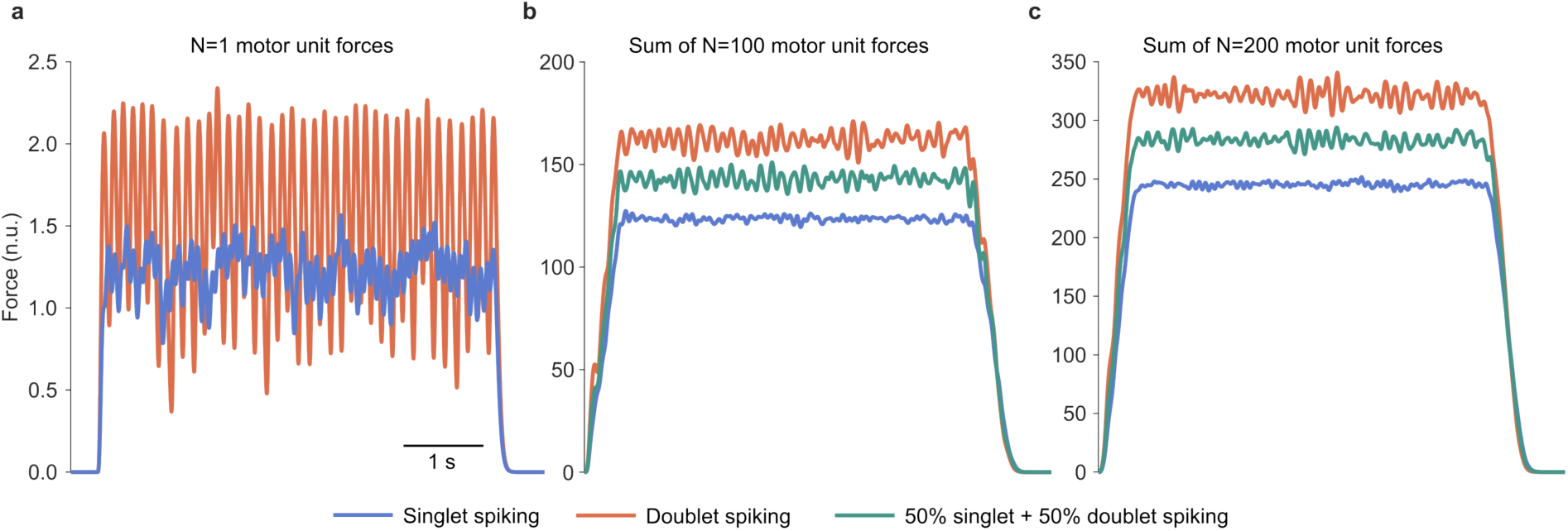
| Repetitive doublets yield a higher mean force at the expense of greater variability. **a**, A simulation of a single motor unit based on the simulations shown in Fig. 7 of the main paper, with motor units exhibiting only repetitive doublets (orange) and only singlet spiking (blue). **b**, Same as in (**a**), but with the sum of 100 motor unit forces, with an additional trace with 50% of the motor units producing singlet spiking and the remaining 50% producing repetitive doublets (green). **c**, Same as in (**b**) but with 200 motor unit traces.

**Extended Data Figure 9.**
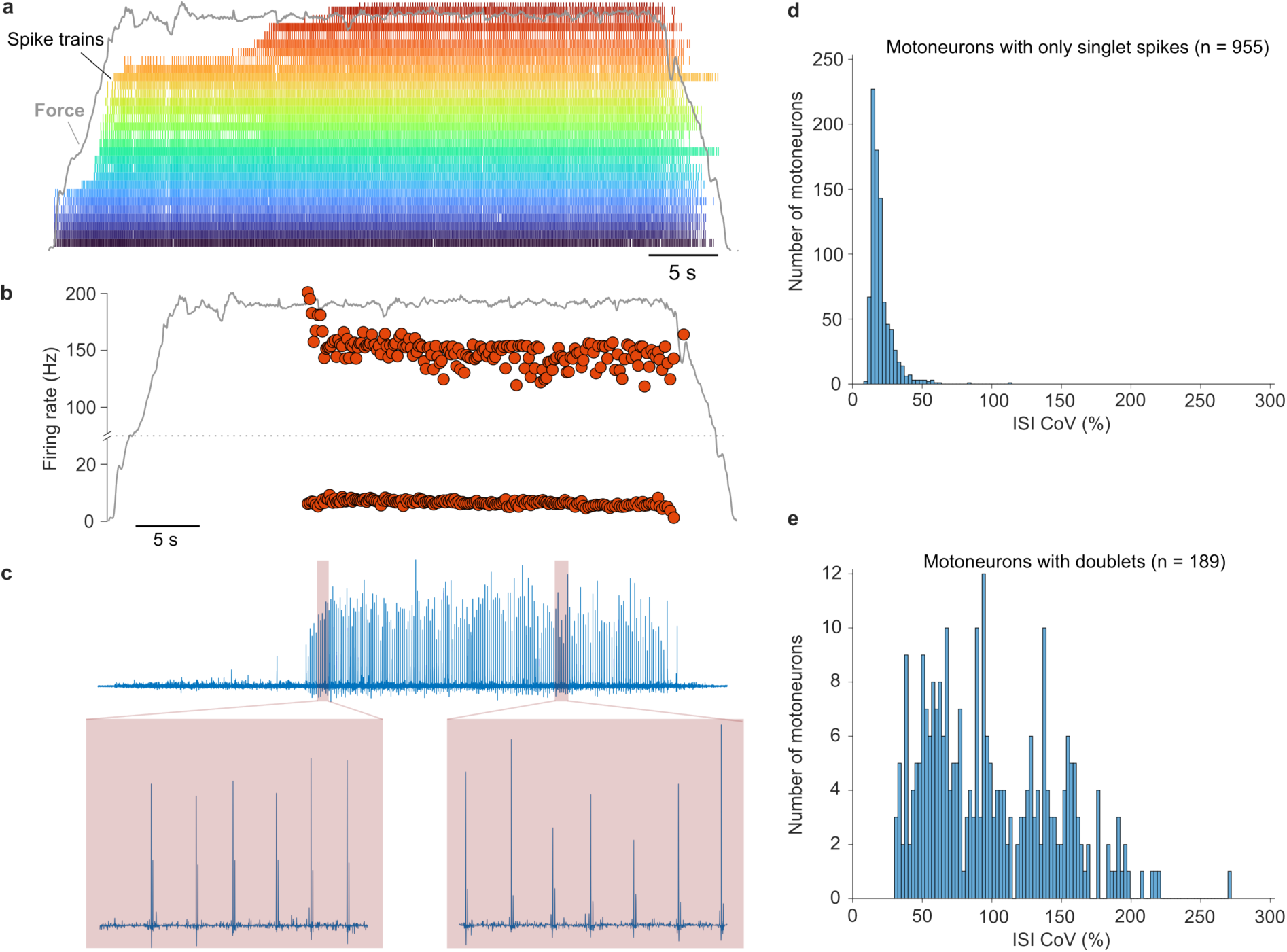
| Motoneuron doublet firings are likely under-reported partly due to methodological biases. **a**, The detected spike trains in subject #13. **b**, One of the last-recruited motoneuron spike trains was detected to produce only repetitive doublets. **c**, Examples of the sources after decomposition show that the second peak of the doublet was initially about half the height of the first peak (spike) and became very small in the later stages, suggesting that only a high signal-to-noise-ratio source will be able to detect such doublets. **d**, The distribution of inter-spike interval coefficient of variation (ISI CoV) for human single-spiking motoneurons (n = 955). **e**, The ISI CoV for human repetitively doublet-spiking motoneurons (n = 189).

## SUPPLEMENTARY METHODS

### S1 Three-compartment conductance-based motoneuron model

The model is a conductance-based, Hodgkin–Huxley-type motoneuron model with three electrotonically coupled compartments representing the dendrite (*V*_d_), soma (*V*_s_) and axon (*V*_a_). The soma is the central compartment lumped with the axon initial segment (AIS), coupled separately to the dendrite and to the axon. The axonal compartment represents the first node of Ranvier. The structure of the model, the kinetic equations of the individual ionic currents and the initial parameter values were adapted from the single-compartment hypoglossal motoneuron model of Purvis & Butera (2005). A subset of maximal conductances was re-scaled from their neonatal-rat reference values to produce firing-rate ranges and spike shapes compatible with human motoneuron recordings; the kinetic expressions of the ionic currents were otherwise left unchanged.

Current was injected into the somatic compartment in all simulations. Unless otherwise stated, voltages are in mV, time in ms, maximal conductances in µS, membrane capacitances in nF, ionic currents in nA and intracellular calcium concentration in arbitrary (dimensionless) units consistent with the original Purvis & Butera (2005) formulation.

#### S1.1 Membrane equations

The three membrane potentials evolve according to

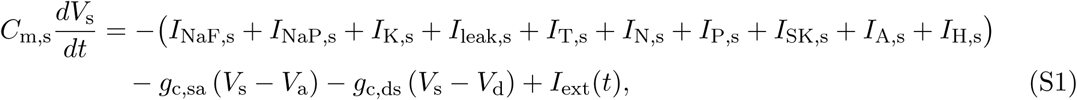

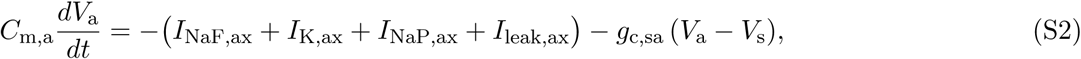

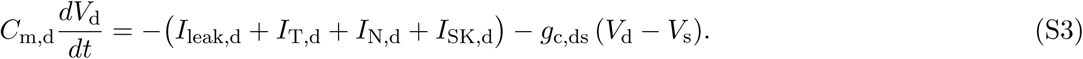

The soma is coupled to the axon through the axial conductance *g*_c,sa_ and to the dendrite through *g*_c,ds_; the dendrite and axon are not directly coupled. The stimulation current *I*_ext_(*t*) is injected into the soma only (Section S2).

#### S1.2 Somatic ionic currents

Each voltage-gated current is the product of a maximal conductance, gating variables raised to integer powers, and an electrochemical driving force:

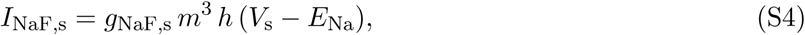

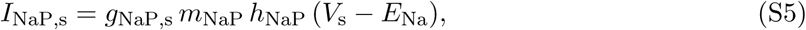

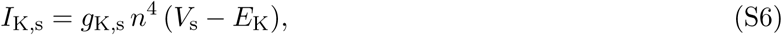

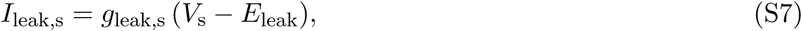

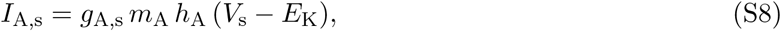

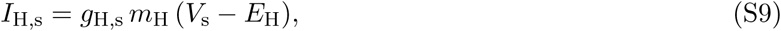

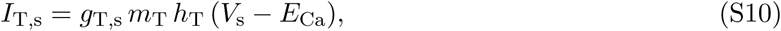

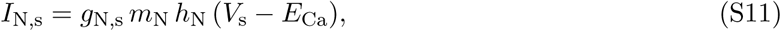

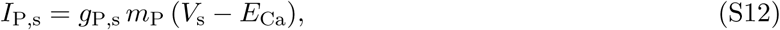

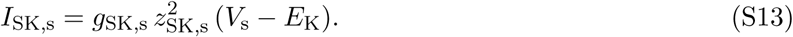

The three somatic calcium currents are of the T-, N- and P-type. The P-type current follows the non-inactivating, high-voltage-activated kinetics reported for hypoglossal motoneurons by Purvis & Butera (2005); we retain the P-type designation rather than relabeling it as L-type, since the kinetics are those of the original model. The total somatic calcium current *I*_Ca,s_ = *I*_T,s_ + *I*_N,s_ + *I*_P,s_ drives the somatic intracellular calcium concentration and hence *I*_SK,s_ through the gating variable *z*_SK,s_.

#### S1.3 Axonal ionic currents

The axonal compartment contains a fast spike-generating sodium current, a delayed-rectifier potassium current, a persistent sodium (NaP) current and a leak (McIntyre et al., 2002):

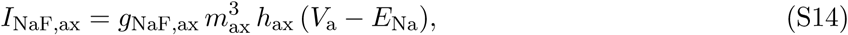

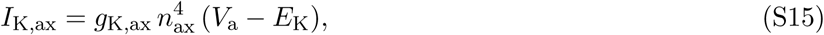

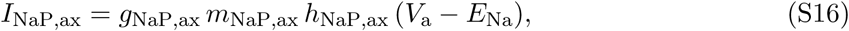

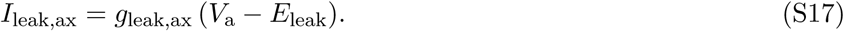

#### S1.4 Dendritic ionic currents

The dendrite is modeled as an active, non-spiking compartment: it does *not* contain fast sodium, delayed-rectifier potassium or A-type potassium currents, in agreement with the low density of action-potential-generating conductances observed in motoneuron dendrites. It contains a leak, T- and N-type calcium currents, and a calcium-activated SK current driven by a dendritic-only calcium pool:

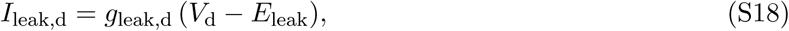

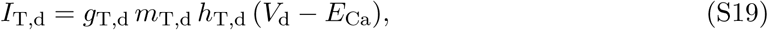

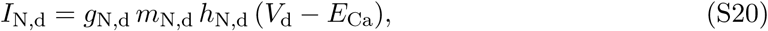

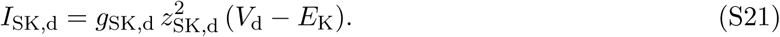

The dendritic P-type calcium conductance was omitted because the spatially averaged dendritic potential never reaches its activation range. The dendritic calcium current that drives the dendritic SK current is therefore *I*_Ca,d_ = *I*_T,d_ + *I*_N,d_. Dendritic calcium dynamics are governed by a dedicated dendritic calcium pool, independent of the somatic pool, so that dendritic SK activation is driven only by local dendritic calcium entry.

#### S1.5 Gating kinetics (soma)

##### Every gating variable *x* obeys

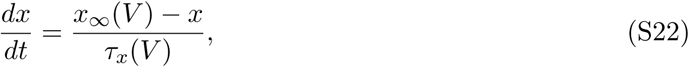

with the following steady-state and time-constant functions, which follow Purvis & Butera (2005). Voltages *V* are in mV and time constants in ms.

##### Fast sodium *I*_NaF_

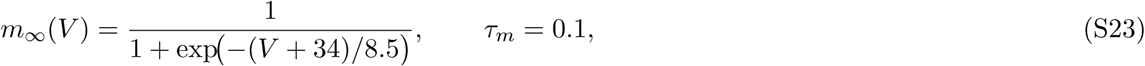

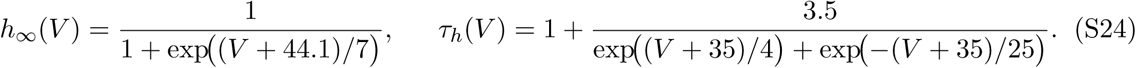

##### Persistent sodium (NaP) *I*_NaP,s_

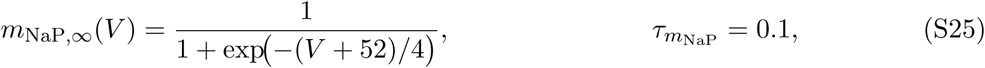

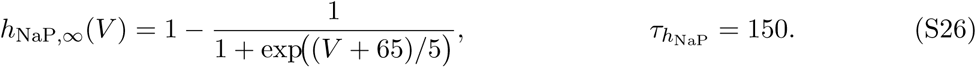

##### Delayed-rectifier potassium *I*_K,s_

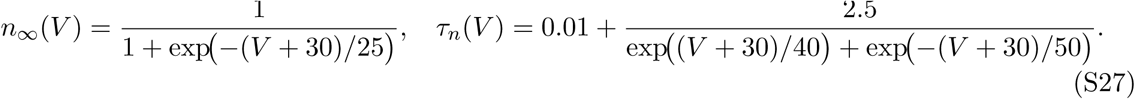

##### T-type calcium *I*_T,s_

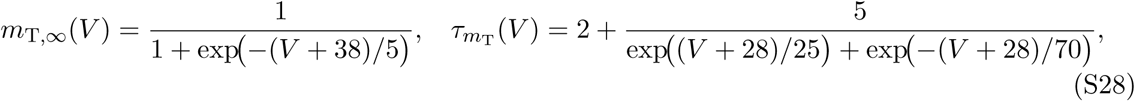

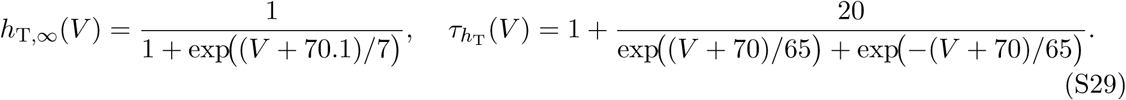

##### N-type calcium *I*_N,s_

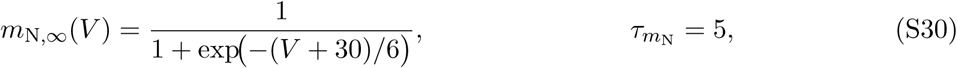

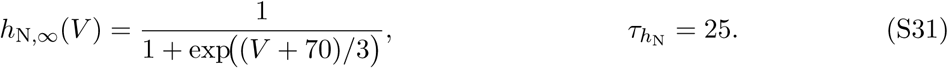

##### P-type calcium *I*_P,s_

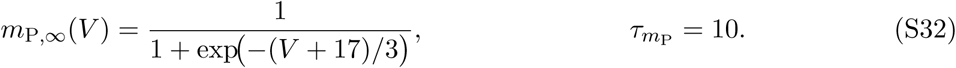

##### A-type potassium *I*_A,s_

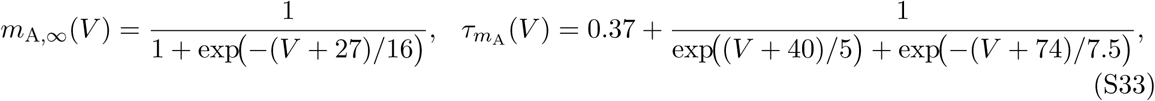

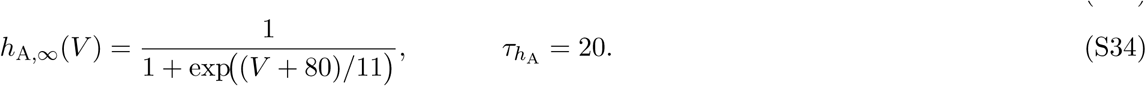

##### HCN current *I*_H,s_

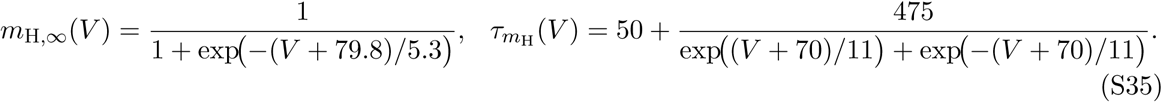

##### Calcium handling and SK activation (soma)

Somatic intracellular calcium is modeled as a first-order buffer driven by the total somatic calcium current:

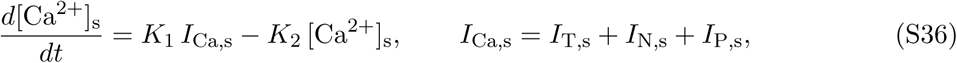

with *K*_1_ = *−*5.3 *×* 10*^−^*^4^ and *K*_2_ = 1.5 *×* 10*^−^*^2^. The SK gating variable is activated by [Ca^2+^]_s_ through a Hill function with half-activation *K_d_* = 0.003 and Hill coefficient 2:

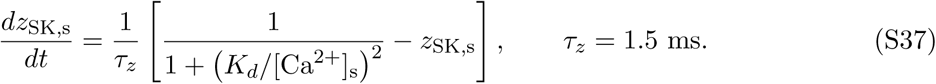

To avoid division by zero during numerical integration, [Ca^2+^]_s_ is floored at 10*^−^*^6^ before evaluation of Eq. (S37).

#### S1.6 Gating kinetics (axon)

The axonal fast sodium and delayed-rectifier potassium currents are described by HH-like kinetics centered around more depolarized voltages and with shorter time constants than their somatic counterparts:

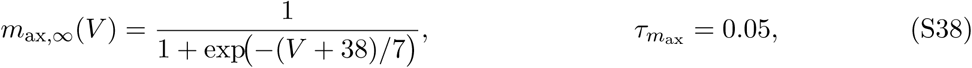

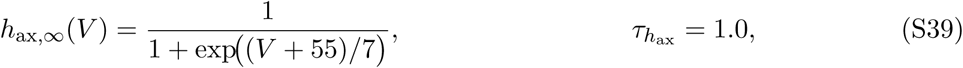

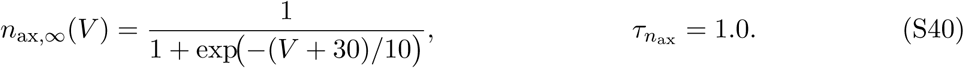

The axonal NaP current has the same Boltzmann form as its somatic counterpart but its own kinetics:

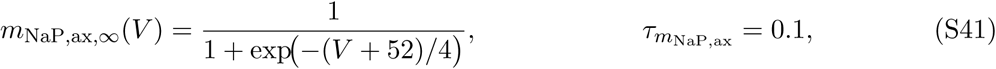

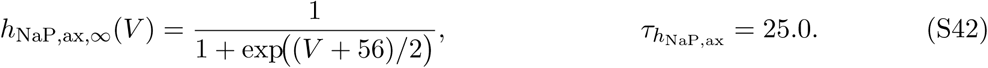

#### S1.7 Gating kinetics (dendrite)

The dendritic T- and N-type calcium channels use the same steady-state and time-constant functions as their somatic counterparts, but are driven by the dendritic membrane potential *V*_d_ and are represented by independent gating variables (*m*_T,d_*, h*_T,d_*, m*_N,d_*, h*_N,d_). The dendritic SK current is driven by a dedicated dendritic calcium pool, which obeys the same buffering equation as the somatic pool but with its own current:

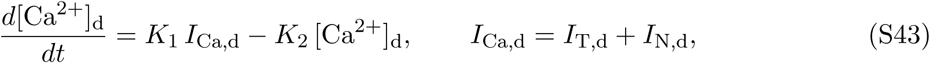

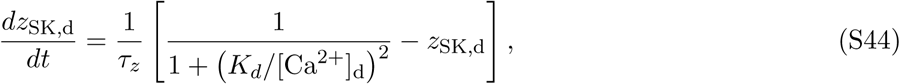

with the same constants *K*_1_, *K*_2_, *K_d_* = 0.003 and *τ_z_* = 1.5 ms as in the soma, and [Ca^2+^]_d_ likewise floored at 10*^−^*^6^.

### S2 Stimulation protocol and post-processing

The injected current *I*_ext_(*t*) is a two-step waveform applied to the soma:

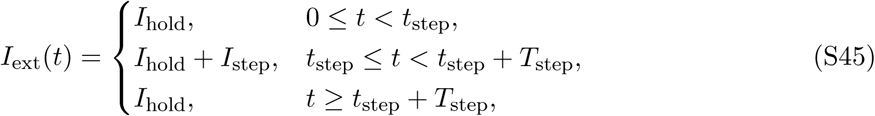

i.e. a long holding current *I*_hold_ followed by a rectangular step of amplitude *I*_step_ and duration *T*_step_. In our reference implementation the pre-stimulus interval was *t*_step_ = 2500 ms, substantially longer than the slowest state variable in the model (*τ_h_*_NaP_ *≈* 150 ms in the soma), so that every state variable reaches steady state well before the step. The step duration *T*_step_ was varied across experiments: a 1 s step was used for the standard firing-pattern and blocking analyzes, and a 10 s step was used to quantify doublet-frequency adaptation (Section S3). The first 1500 ms of each simulation were discarded from all analyses.

### S3 Quantification of doublet-frequency adaptation

To quantify the doublet-frequency adaptation during repetitive doublet firing, we analyzed the somatic voltage trace *V*_s_(*t*) produced by a 10 s step current (Section S2). Spikes were detected from *V*_s_(*t*) using a local-maximum criterion with a voltage threshold of 0 mV and a minimum inter-peak interval of 2 ms. For each pair of consecutive spikes we computed the instantaneous firing rate *f_i_* = 1000*/*(*t_i_*_+1_ *− t_i_*), where *t_i_* is the time of the *i*-th spike in milliseconds. To isolate the intra-doublet (high-frequency) intervals, only intervals with *f_i_ ≥* 100 Hz were retained; these constitute the short interval within each doublet, as opposed to the long inter-doublet intervals. The first retained intra-doublet interval was excluded, because the first doublet is shaped by the onset of the injected drive. The retained intra-doublet frequencies *f_k_*, as a function of the time *t_k_*of each doublet (taken as the time of the first spike of the doublet), were fitted with a decaying exponential,

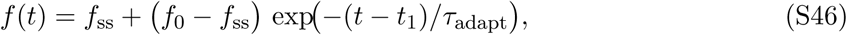

where *t*_1_ is the time of the first retained doublet, *f*_ss_ is the steady-state intra-doublet frequency, *f*_0_ is the (extrapolated) initial frequency, and *τ*_adapt_ is the adaptation time constant (in ms). The fit was performed by nonlinear least squares. We report *f*_ss_ and *τ*_adapt_ as functions of the somatic NaP inactivation time constant, which was varied by modifying *τ_h_*_NaP_.

### S4 Numerical integration

The coupled ODEs (S1)–(S3), together with the gating and calcium equations, form a system of state variables. The system was integrated with a fixed-step, classical fourth-order Runge–Kutta (RK4) scheme:

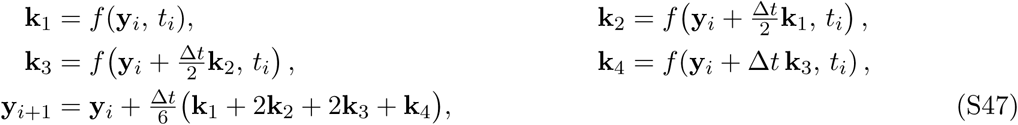

with integration step Δ*t* = 1 µs, implemented in MATLAB (MathWorks, Natick, MA, USA). After each step, gating variables were clipped to [0, 1] and intracellular calcium concentrations to [0*, ∞*) to suppress rare numerical excursions outside the biophysically meaningful range.

Initial conditions for all state variables are listed in Supplementary Table S3; all three compartments are initialized at *−*73.8 mV. After the long pre-stimulus interval these initial values are effectively overwritten by the steady state of Eq. (S45) with *I*_ext_ = *I*_hold_, so they have no effect on the reported analyses.

### S5 Parameters

Maximal conductances and calcium-handling parameters were taken from Purvis & Butera (2005) as a starting point, and a subset was re-scaled manually so that the model reproduced the firing-rate ranges and firing patterns (single spikes, repetitive firing, initial doublets, and repetitive doublets) observed in human motoneuron recordings. The potassium reversal potential *E*_K_ was set to *−*90 mV for the initial-doublet regime and to *−*85 mV for the repetitive-doublet regime (main text). Nominal values are listed in Table S1 and Table S2; the amplitudes of the holding and step currents for each simulated condition are given in the corresponding figure captions.

**Table S1:** Reversal potentials and passive properties.

| Symbol | Value | Description |
| --- | --- | --- |
| $E_{Na}$ | +60 mV | $Na^+$ reversal potential |
| $E_K$ | $-90 / -85$ mV | $K^+$ reversal potential (initial / repetitive regime) |
| $E_{Ca}$ | +50 mV | $Ca^{2+}$ reversal potential |
| $E_H$ | $-38.8$ mV | HCN reversal potential |
| $E_{leak}$ | $-70$ mV | Leak reversal potential |

---

**Table S2:** Maximal conductances, capacitances and calcium-handling parameters of the three-compartment model. Values correspond to the nominal (control) configuration.

| Symbol | Value | Description |
| --- | --- | --- |
| *Soma (including AIS)* |  |  |
| $g_{\text{NaF},s}$ | 1.3 | Fast $\text{Na}^+$ |
| $g_{\text{K},s}$ | 1.5 | Delayed-rectifier $\text{K}^+$ |
| $g_{\text{leak},s}$ | 0.008 | Leak |
| $g_{\text{A},s}$ | 1.1 | A-type $\text{K}^+$ |
| $g_{\text{H},s}$ | 0.005 | HCN |
| $g_{\text{NaP},s}$ | 0.03 | Persistent $\text{Na}^+$ |
| $g_{\text{SK},s}$ | 0.30 | SK |
| $g_{\text{T},s}$ | 0.05 | T-type $\text{Ca}^{2+}$ |
| $g_{\text{N},s}$ | 0.02 | N-type $\text{Ca}^{2+}$ |
| $g_{\text{P},s}$ | 0.02 | P-type $\text{Ca}^{2+}$ |
| $C_{\text{m},s}$ | 0.08 | Capacitance |
| *Axon (first node of Ranvier)* |  |  |
| $g_{\text{NaF},\text{ax}}$ | 0.20 | Fast $\text{Na}^+$ |
| $g_{\text{K},\text{ax}}$ | 0.60 | Delayed-rectifier $\text{K}^+$ |
| $g_{\text{NaP},\text{ax}}$ | 0.04 | Persistent $\text{Na}^+$ |
| $g_{\text{leak},\text{ax}}$ | 0.004 | Leak |
| $C_{\text{m},\text{a}}$ | 0.040 | Capacitance |
| *Dendrite* |  |  |
| $g_{\text{SK},\text{d}}$ | 0.30 | SK |
| $g_{\text{leak},\text{d}}$ | 0.025 | Leak |
| $g_{\text{T},\text{d}}$ | 0.08 | T-type $\text{Ca}^{2+}$ |
| $g_{\text{N},\text{d}}$ | 0.15 | N-type $\text{Ca}^{2+}$ |
| $C_{\text{m},\text{d}}$ | 0.20 | Capacitance |
| *Coupling and calcium handling* |  |  |
| $g_{\text{c},\text{sa}}$ | 0.20 | Soma–axon coupling |
| $g_{\text{c},\text{ds}}$ | 0.07 | Soma–dendrite coupling |
| $K_1$ | $-5.3 \times 10^{-4}$ | $\text{Ca}^{2+}$ influx coupling (both pools) |
| $K_2$ | $1.5 \times 10^{-2}$ | $\text{Ca}^{2+}$ extrusion rate (both pools) |
| $[\text{Ca}^{2+}]_{\text{init}}$ | 0.10 | Initial $\text{Ca}^{2+}$ (both pools) |

**Table S3:** Initial conditions for all state variables. After the long pre-stimulus interval these values have no effect on the reported simulations.

| Variable | Initial value | Compartment |
| --- | --- | --- |
| $V_d, V_s, V_a$ | $-73.8$ mV | all |
| $m$ | 0.015 | soma |
| $h$ | 0.981 | soma |
| $m_{\text{NaP},s}$ | 0.002 | soma |
| $h_{\text{NaP},s}$ | 0.797 | soma |
| $n$ | 0.158 | soma |
| $m_{\text{T},s}$ | 0.001 | soma |
| $h_{\text{T},s}$ | 0.562 | soma |
| $m_{\text{P},s}$ | 0 | soma |
| $m_{\text{N},s}$ | 0.001 | soma |
| $h_{\text{N},s}$ | 0.649 | soma |
| $z_{\text{SK},s}$ | 0 | soma |
| $m_{\text{A},s}$ | 0.057 | soma |
| $h_{\text{A},s}$ | 0.287 | soma |
| $m_{\text{H},s}$ | 0.182 | soma |
| $[\text{Ca}^{2+}]_s$ | $[\text{Ca}^{2+}]_{\text{init}}$ | soma |
| $m_{\text{T},d}$ | 0.001 | dendrite |
| $h_{\text{T},d}$ | 0.562 | dendrite |
| $m_{\text{N},d}$ | 0.001 | dendrite |
| $h_{\text{N},d}$ | 0.649 | dendrite |
| $z_{\text{SK},d}$ | 0 | dendrite |
| $[\text{Ca}^{2+}]_d$ | $[\text{Ca}^{2+}]_{\text{init}}$ | dendrite |
| $m_{\text{NaP},\text{ax}}$ | 0 | axon |
| $h_{\text{NaP},\text{ax}}$ | 1 | axon |
| $m_{\text{ax}}$ | 0.01 | axon |
| $h_{\text{ax}}$ | 0.98 | axon |
| $n_{\text{ax}}$ | 0.02 | axon |

